# Evaluating generalizability of artificial intelligence models for molecular datasets

**DOI:** 10.1101/2024.02.25.581982

**Authors:** Yasha Ektefaie, Andrew Shen, Daria Bykova, Maximillian Marin, Marinka Zitnik, Maha Farhat

**Author notes:** Corresponding authors: yasha, maha. Contributed equally (Marinka Zitnik, Maha Farhat).

## Abstract

Deep learning has made rapid advances in modeling molecular sequencing data. Despite achieving high performance on benchmarks, it remains unclear to what extent deep learning models learn general principles and generalize to previously unseen sequences. Benchmarks traditionally interrogate model generalizability by generating metadata based (MB) or sequence-similarity based (SB) train and test splits of input data before assessing model performance. Here, we show that this approach mischaracterizes model generalizability by failing to consider the full spectrum of cross-split overlap, *i.e.*, similarity between train and test splits. We introduce SPECTRA, a spectral framework for comprehensive model evaluation. For a given model and input data, SPECTRA plots model performance as a function of decreasing cross-split overlap and reports the area under this curve as a measure of generalizability. We apply SPECTRA to 18 sequencing datasets with associated phenotypes ranging from antibiotic resistance in tuberculosis to protein-ligand binding to evaluate the generalizability of 19 state-of-the-art deep learning models, including large language models, graph neural networks, diffusion models, and convolutional neural networks. We show that SB and MB splits provide an incomplete assessment of model generalizability. With SPECTRA, we find as cross-split overlap decreases, deep learning models consistently exhibit a reduction in performance in a task- and model-dependent manner. Although no model consistently achieved the highest performance across all tasks, we show that deep learning models can generalize to previously unseen sequences on specific tasks. SPECTRA paves the way toward a better understanding of how foundation models generalize in biology.

## Main

Understanding generalizability – how well a machine learning model performs on unseen data – is a fundamental challenge for the broad use of computation in biological discovery. In living cells, information flows from DNA to RNA to protein and dictates cell phenotypes. To model phenotypes, deep learning models are trained to predict biological relationships between and within sequences and the phenotype. This approach has been successfully implemented through a broad array of machine learning models, including convolutional neural networks [1–4], recurrent neural networks [5–7], graph neural networks [8–11], and large language models [12–16]. However, model evaluation is challenging because (1) available molecular sequencing data often capture a small fraction of all possible sequences due to the time and cost of generating sequencing data [17, 18]. (2) Sequences evolve and acquire new mutations over time that are not present in existing datasets. This results in differences in the distribution of sequences and their aggregate properties between datasets, known as distribution shifts that can lead to degradation of model performance [19–21]. Although distributional shifts are a well-recognized challenge in machine learning more generally [22, 23], they are less well characterized in biology due to the lack of approaches that measure model performance in the context of distribution shifts. Though numerous benchmarks have been developed to assess model performance across datasets [16, 24–27], there are still large gaps between model performance during benchmarking and real-world use [28–32] (Figure 1a). This gap in assessing generalizability must be addressed before machine learning models can be broadly used in biology.

**Figure 1:**
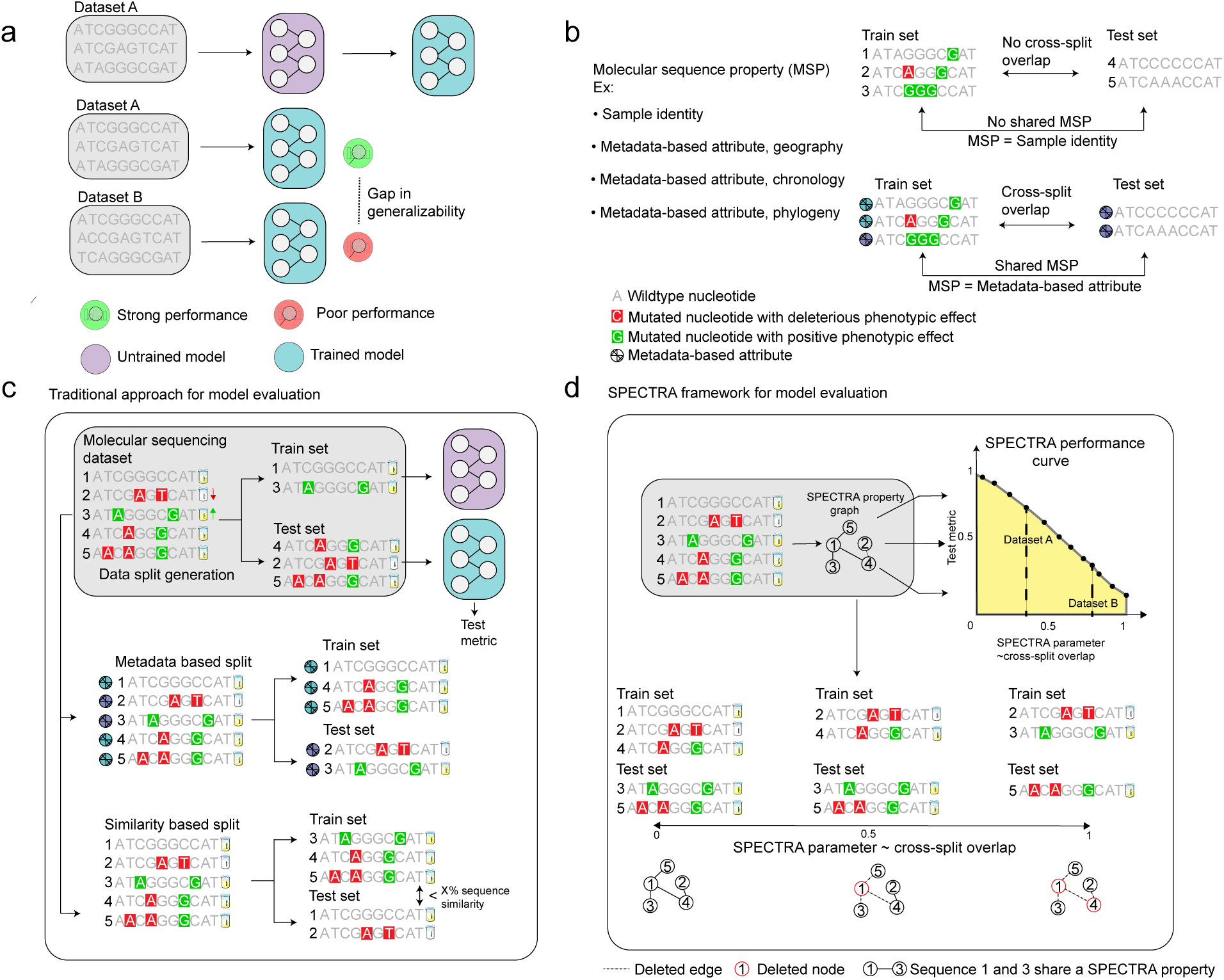
The SPECTRA framework for model evaluation (SPECTRA). **(a)** Machine learning (ML) models for molecular sequencing data struggle to generalize across datasets. **(b)** Every train-test split partitions samples based on a chosen molecular sequence property. Cross-split overlap exists between train and test sets when samples share molecular sequence properties. Shown are two examples of train-test splits. In the first, samples are split based on their identity, shown by the sample number, and there is no cross-split overlap as no two samples share identity across the train and test set. In the second, samples are split based on metadata-based attributes with cross-split overlap as two samples share an attribute across the train and test set. **(c)** Traditional approaches for model evaluation create train-test splits either based on metadata-based attributes (metadata based split) or sequence similarity (similarity based split). **(d)** The SPECTRA or spectral framework for model evaluation generates train-test splits with a spectrum of cross-split overlap. It does so by constructing a spectral property graph where nodes are samples and edges are between samples that share a spectral property. The spectral property is a molecular sequence property which influences model generalizability. It then iteratively deletes nodes and edges based on the spectral parameter, an internal parameter, to generate train and test sets. After training and evaluating an input model to each generated split, SPECTRA generates a spectral performance curve which plots model test performance versus the spectral parameter, an internal parameter that scales with cross-split overlap. The spectral performance curve shows the gap in generalizability depicted in panel a is due to differing levels of cross-split overlap between splits generated in dataset A and those of dataset B. The area under the spectral performance curve (highlighted in yellow) summarizes model performance across all levels of cross-split overlap and is a new metric for model generalizability. Note: The use of the word spectral here refers only to the framework for model evaluation and should not be confused with other uses of the term in matrix analysis.

While useful, the central shortcoming of existing benchmarks is the approach to model evaluation. Existing methods for model evaluation split input molecular sequencing datasets into train and test sets in metadata-based (MB) or similarity-based (SB) splits (Figure 1c). MB splits ensure certain metadata properties are not shared across splits. One example is a temporal split of Covid-19 viral sequences in which a vaccine escape model is trained on sequences collected before a specific time and tested on sequences evolved after that time [33–35]. A random split is also an MB split where the metadata property is sample identity. SB splits ensure no two samples across splits share sequence similarity beyond a pre-defined threshold, with the exact threshold being problem-specific [36–40]. However, as we show in this study, MB splits cannot guarantee that high performance on the test set will transfer to a new molecular sequencing dataset. This is because metadata-based partitioning does not control sequence similarity between data subsets. Model generalizability can be overestimated when training sequences are more similar to sequences in the test set than to sequences in a new dataset. In SB splits, sequence similarity can be controlled. However, the generalizability of the model at similarity thresholds different from a handful of those tested during model benchmarking remains unknown, resulting in an incomplete evaluation of the model. Further, SB splits rely on limited summary metrics such as sequence distance to quantify similarity between sequences that may not capture the full range of similarities. This lack of understanding about model generalizability can lead to several significant issues. Firstly, there could be catastrophic degradation of model performance on new datasets, which means that predictions made on unseen data may be highly inaccurate. This inaccuracy can mislead biological research, causing wasted resources on false leads or overlooking potential discoveries. Secondly, models that perform well on training data but poorly on unseen data can contribute to overfitting, where the model captures noise rather than the underlying biological processes. This overfitting can severely limit the applicability of computational tools in novel or broader biological contexts, potentially leading to erroneous conclusions about biological mechanisms and phenotypes.

Here, we introduce the spectral framework for model evaluation (SPECTRA)^1^, a framework for evaluating generalizability of machine learning models for molecular sequences. Given a model, a molecular sequencing dataset, and a spectral property definition, SPECTRA generates a series of train-test splits with decreasing overlap, i.e., a spectrum of train-test splits. SPECTRA then plots the model’s performance as a function of cross-split overlap (Figure 1b,d) generating a spectral performance curve (SPC). We propose the area under this curve (AUSPC), as a new more comprehensive metric for model generalizability. We apply SPECTRA to 18 molecular sequencing datasets from three prominent benchmarks (PEER [25], ProteinGym [16], TAPE [24]) and find (1) existing SB and MB splits have large amounts of cross-split overlap, (2) SPECTRA generates splits with the same level of cross-split overlap than existing SB and MB splits, and (3) existing SB and MB splits represent single points in the SPC, leaving the rest of the SPC uncharacterized. To demonstrate the need to characterize model SPCs, we apply SPECTRA to eleven state-of-the-art machine learning models including pretrained and finetuned large language models (LLM), convolutional neural networks (CNN), graph neural networks (GNN), variational autoencoders (VAE), and diffusion generative models. We find that none of the machine learning models achieves a high AUSPC across all tested tasks. We show that examination of the SPC can help identify unconsidered spectral properties that influence model generalizability in molecular sequencing datasets. By applying SPECTRA to pretrained protein language models, we demonstrate how SPECTRA can be used to evaluate foundation models in biology. SPECTRA is a novel paradigm for model evaluation given its ability to more comprehensively characterize model generalizability and uncover short-comings of existing machine learning models. These capabilities will inform the next generation of machine learning models for molecular sequencing data.

## Results

### Overview of the spectral framework for model evaluation (SPECTRA)

In contrast to the traditional approach to machine learning model evaluation using meta-data based (MB) and similarity based (SB) splits, the spectral framework (SPECTRA) provides a more comprehensive overview of model performance by examining a model’s spectral performance curve for a given molecular sequencing dataset. The approach relies on focusing on one or more characteristics of input molecular sequences or molecular sequence properties (MSP) (*e.g.* GC content of a gene). We define a spectral property (SP) as a MSP expected to affect model generalizability for a specific task (*e.g.* 3D protein structure for protein binding prediction). The definition of the spectral property is task-specific and, together with the molecular sequence dataset and model, are the only inputs to SPECTRA (Figure 1d). First, SPECTRA compares the spectral property for all pairs of sequences in the dataset, identifying pairs that share the spectral property. The procedure is used to construct a spectral property graph from which adaptive train-test splits are generated with decreasing cross-split overlap or the proportion of samples in the test set that share a SP with the train. SPECTRA controls cross-split overlap by ranging an internal spectral parameter (SP) from SP = 0 to SP = 1 for maximal and minimal cross-split overlap, respectively. Lastly, the model is trained and tested on each split to generate a plot of the model’s performance against the spectral parameter, the model’s spectral performance curve (SPC), for the molecular sequencing dataset. The area under the spectral performance curve (AUSPC) summarizes model performance across all levels of cross-split overlap and can be used to compare model generalizability to other models within and across tasks.

### SPECTRA expands on traditional approaches to model evaluation

We hypothesize that SB and MB splits used in existing machine learning benchmarks represent individual points on the SPC and that SPECTRA provides a more comprehensive view of model generalization compared to prevailing dataset splits. To validate this proposition, we applied SPECTRA to six molecular sequencing datasets from two widely-used protein sequence benchmarks, TAPE [24] and PEER [25], as well as a protein structural biology benchmark, PDBBind [41], and a protein mutational benchmark, ProteinGym [16] (Section 2.4-2.9 Methods). First, we were interested in what specific points on SPC each dataset split in these benchmarks corresponds to. Specifically, we calculated the cross-split overlap in the MB and SB splits implemented in these benchmarks and identified the spectral parameter that generates splits with similar cross-split overlap. For example, our analyses using SPECTRA revealed that the MB family split from the remote homology dataset in TAPE had a cross-split overlap of 97% and can be instantiated at SP = 0.025. In another example, the temporal-based MB split of PDBBind used to train the Equibind model had a cross-split overlap of 76% and was generated at SP = 0.55 [35] (Figure 2a).

**Figure 2:**
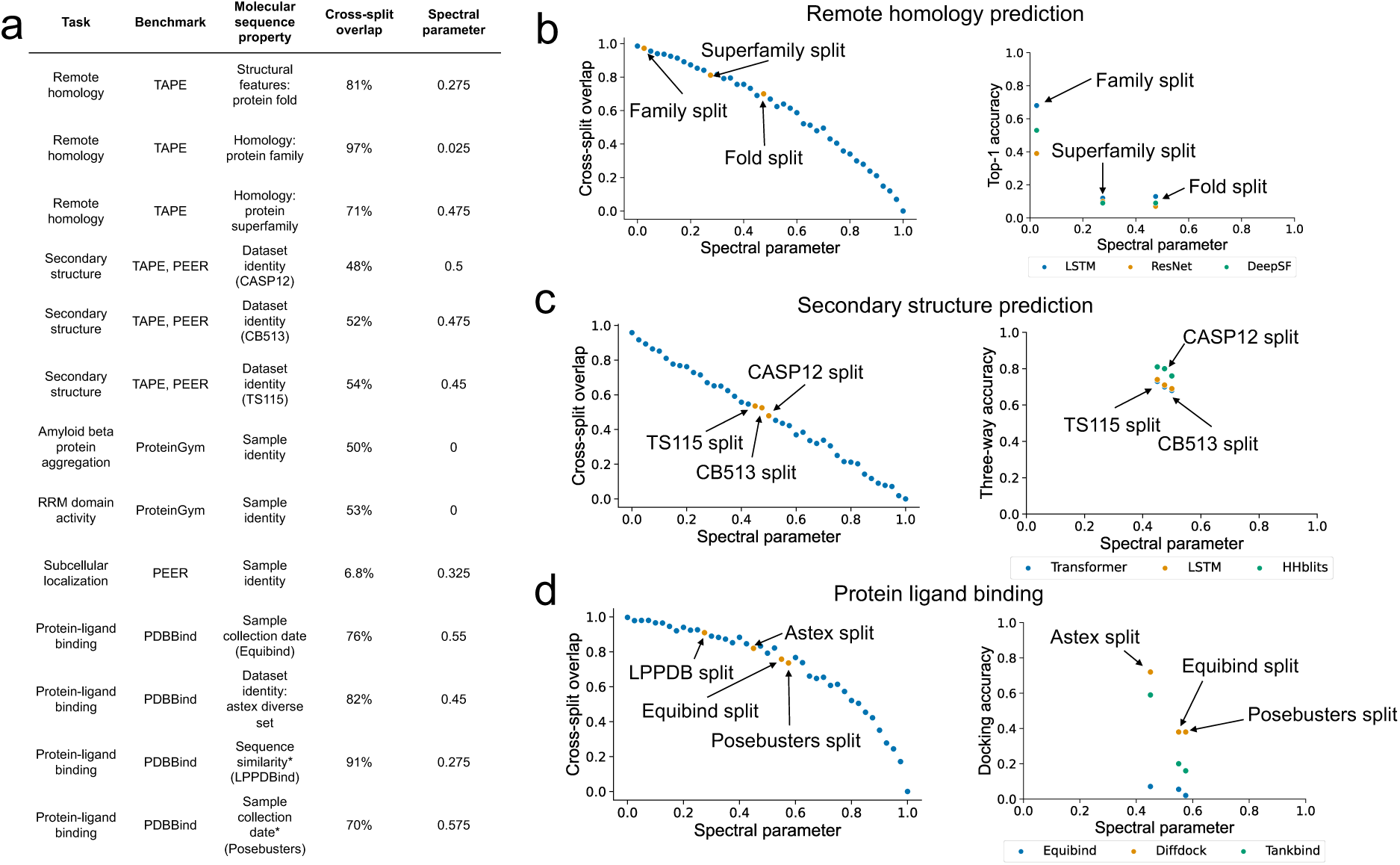
Applying SPECTRA to existing molecular sequencing benchmarks. **(a)** Results of applying SPECTRA on datasets from the PEER, TAPE, ProteinGym, and PDBBind benchmarks. For every dataset we report the corresponding task and benchmark, the molecular sequence property that was used to split the dataset into train and test, cross-split overlap, and the spectral parameter in SPECTRA that generated a split with similar levels of cross-split overlap. If a specific name is used to refer to the split in literature, the name is provided in paranthesis. (*Posebusters and LPPDBind use multiple manually defined quality control criteria to create splits) **(b)** (Left) Cross-split overlap as spectral parameter increases in SPECTRA for the TAPE remote homology benchmark. Labeled are the points on the curve where the family, superfamily, and fold splits have similar levels of cross-split overlap. (Right) A partial spectral performance curve for a LSTM [42], ResNet [72], and DeepSF [43] model. **(c)** (Left) Cross-split overlap as spectral parameter increases in SPECTRA for the TAPE secondary structure benchmark. Labeled are the points on the curve where the CASP12 [73], TS115 [74], and CB513 [75] splits have similar levels of cross-split similarity. (Right) A partial spectral performance curve for a Transformer [49], LSTM [42] and HHblits [76] model. **(d)** (Left) Cross-split overlap as spectral parameter increases in SPECTRA for the PDBBind benchmark. Labeled are the points on the curve where the LPPDBind [67], Astex diverse set [77], Equibind [35] and Posebusters [66] splits have similar levels of cross-split similarity. (Right) A partial spectral performance curve for Equibind [35], Diffdock [78], and Tankbind [79] models.

We also found that in general model performance decreased with decreasing cross-split overlap. The accuracy of long short-term memory (LSTM) [42] and convolutional neural network (CNN) [43] models decreased by 50% between family and superfamily splits in the TAPE benchmark’s remote homology dataset. The cross-split overlap was lower for the superfamily (71%) compared to the family split (97%) (Figure 2b). We observed a similar pattern when using SPECTRA to study models for predicting secondary protein structure (Figure 2c) and protein-ligand binding affinity (Figure 2d). This indicates that existing molecular benchmarks capture only a few points on the spectral performance curve providing a myopic assessment of model generalizability. Further, the observation that model performance diminishes when cross-split overlap decreases identifies an overestimation of model performance by existing benchmarks which would lead to suboptimal performance when the models are implemented in the real world.

### SPECTRA reveals generalization gaps in state-of-the-art molecular machine learning models

To demonstrate the utility of SPECTRA in characterizing the full model spectral performance curve, we evaluate six models in five molecular sequencing datasets. Specifically, we generate SPECTRA splits for each dataset, train and test models on each split, generate the spectral performance curve and calculate the area under the spectral curve for each model.

Our data spans three diverse problems: antibiotic resistance in *M. tuberculosis* (TB) [1], vaccine escape in severe acute respiratory syndrome coronavirus 2 (SARS-CoV-2) [44], and fluorescence prediction in the green fluorescent protein (GFP) of *Aequorea victoria* [45]. Antibiotic resistance in TB is a whole organism phenotype where inputs are the nucleic acid sequences of genes and non-coding regions causally linked to resistance to the specific drug. The output is resistance binary phenotype determined using a culture-based assay. We consider TB resistance to the antibiotics rifampicin (RIF), isoniazid (INH), and pyrazinamide (PZA). For fluorescence prediction of *Aequorea victoria*, we rely on the amino acid sequence of GFP protein and its variants. Vaccine escape in SARS-CoV-2 maps mutations in the receptor binding domain of the spike protein to a continuous value that represents antibody escape (Section 2.1-2.3 Methods).

We evaluated six approaches to modeling phenotype from molecular sequence data. We generated spectral performance curves for a logistic regression model, a convolutional neural network (CNN [1]), a pretrained (GearNet [8]) and fine-tuned structure-based graph neural network (GearNet-Finetuned), a pre-trained (Evolutionary Scale Modeling or ESM2 [12]), and a fine-tuned large language model (ESM2-Finetuned), a multiple sequence alignment-based generative model (GM) (evolutionary model of variant effect prediction - EVE [33]) and an alignment-free GM (SeqDesign [46]) (Section 3.1 Methods).

As was seen for the protein benchmarks above, all evaluated models demonstrate a decrease in performance as cross-split overlap decreases (Figure 3a). Logistic regression decreases performance from an AUC greater than 0.9 for RIF and INH resistance prediction to 0.5 (RIF AUSPC = 0.86, INH AUSPC = 0.74). ESM2-Finetuned decreases in performance for GFP from a Spearman’s rank correlation greater than 0.9 to one less than 0.4. Although all models demonstrate decreased performance with decreased cross-split overlap, some models continue to perform well at minimum cross-split overlap (SP = 1). In RIF and PZA, ESM2, ESM2-Finetuned, and CNN maintain AUCs greater than 0.7 at SP = 1 (Figure 3a). No single model outperforms others across all tasks by AUSPC (Figure 3b). CNN has an AUSPC greater than 0.6 across all tasks but is surpassed by ESM2-Finetuned and ESM2 in RIF (Figure 3b).

**Figure 3:**
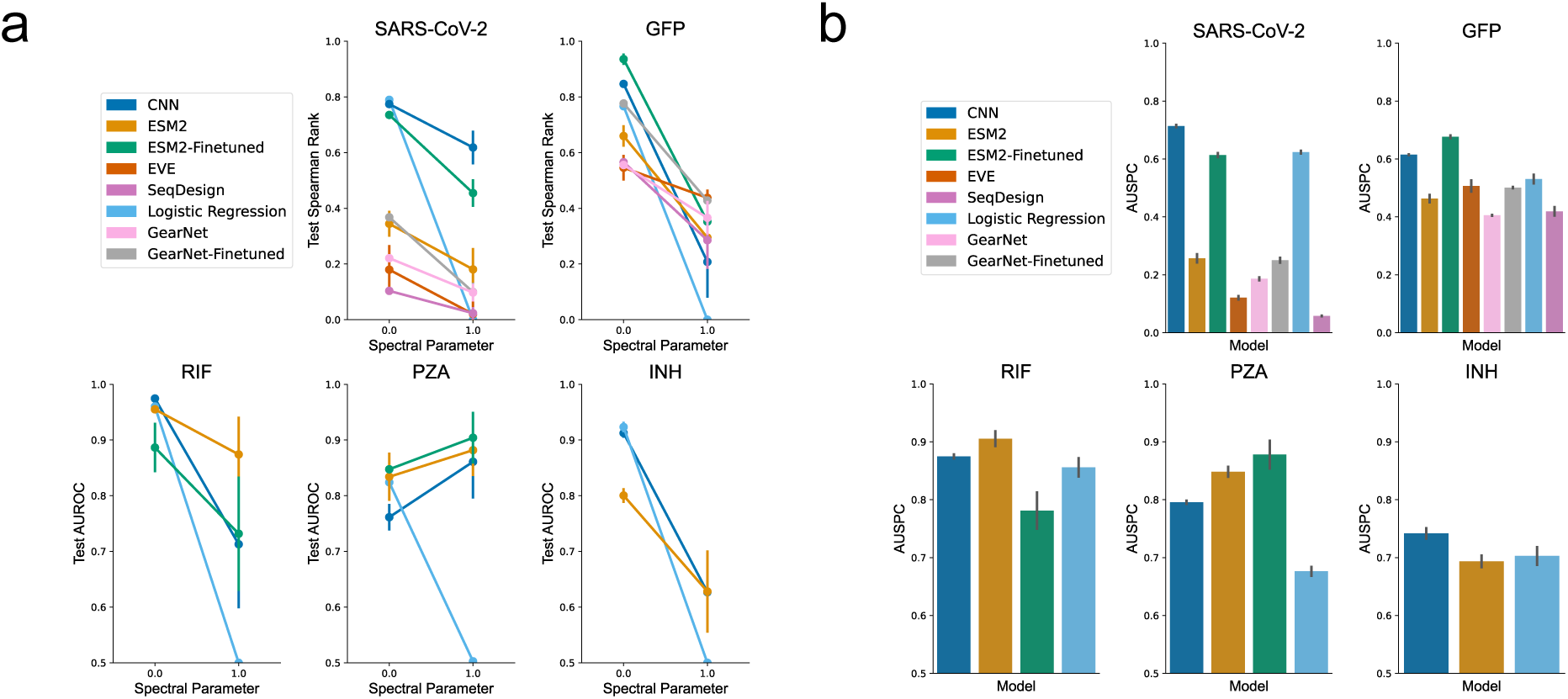
Application of SPECTRA to eight machine learning models across five molecular sequencing datasets. **a.** Test metric change between spectral parameter of 0 (maximum cross-split overlap) and 1 (no cross split overlap) across all tested models and tasks. **b.** Average AUSPC across all models and tasks. Y-axis begins at random AUSPC.

### SPECTRA uncovers spectral properties learned by machine learning models

Every spectral performance curve represents a hypothesis that model performance will vary as a function of the defined spectral property. Any additional deviation of model performance from the expected performance under the defined SPC may indicate the presence of unconsidered spectral properties. By identifying these spectral properties, we can better understand what models learn from molecular sequencing datasets and identify shortcomings of existing models.

CNN model performance demonstrates a high variance across splits with the same spectral parameter suggesting the presence of an unconsidered spectral property (Figure 4a). Three splits in RIF at SP = 0.9, SP = 0.95, and SP = 1.0 have an AUROC standard deviation (and total performance decrease) of 0.09 (26%), 0.10 (31%), and 0.08 (23%) respectively (Figure 4a). RIF resistance is mainly caused by missense mutations in an 81 base pair region of the RNA polymerase beta-subunit (*rpoB*) gene, the rifampicin resistance determining region (RRDR) (Figure 4b). This region spans the active DNA binding site of the polymerase and binds the drug rifampicin resulting in inhibition of transcription. [47, 48]. Models able to learn the association of RRDR to resistance most comprehensively will achieve high test performance. We hypothesize that as SP increases, fewer RRDR mutations are observed in training and the genetic distance between observed RRDR mutations in the train and test increases. On such splits, models may associate partial regions of the RRDR to resistance that do not align with RRDR regions in the test, resulting in poor generalizability.

**Figure 4:**
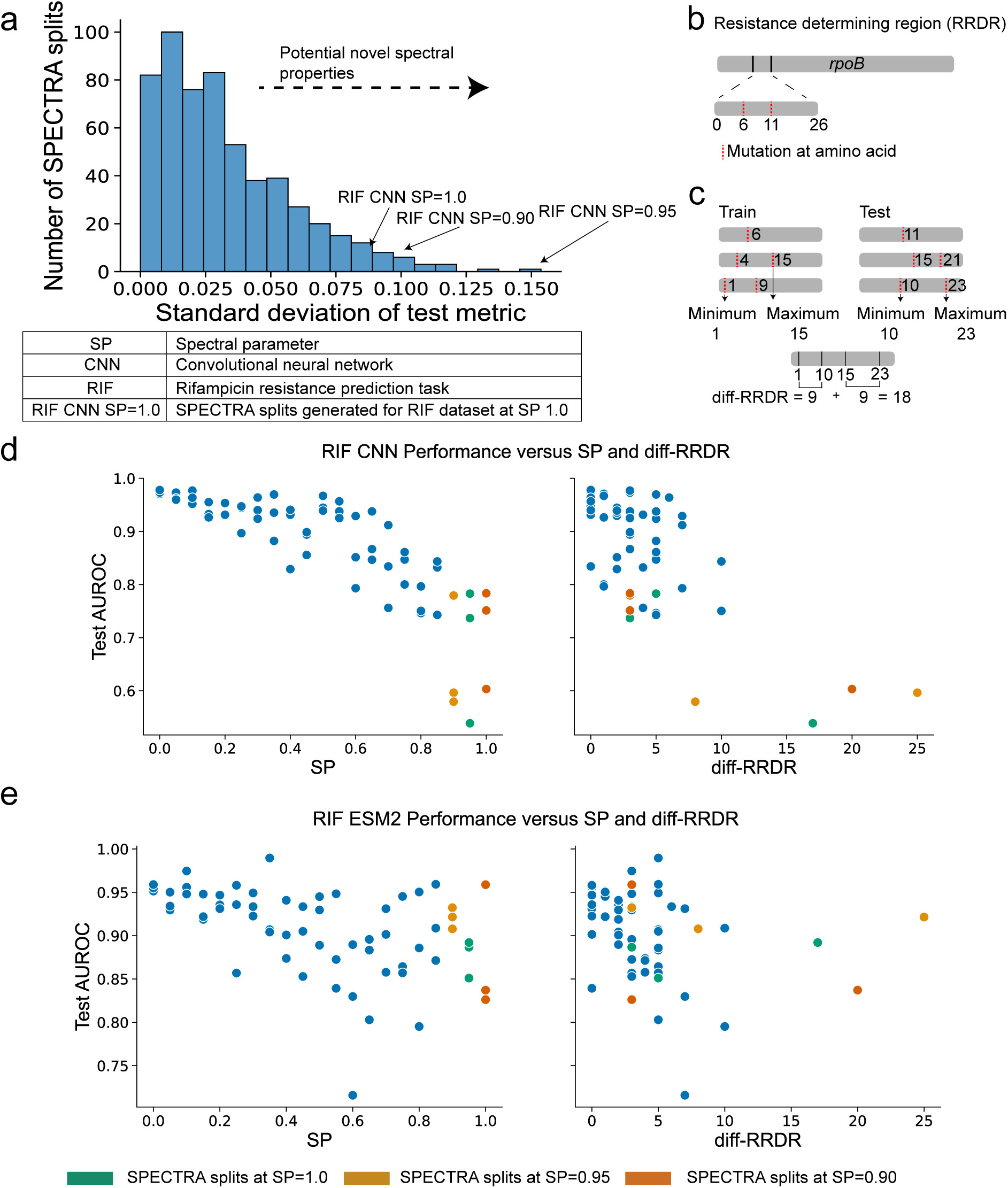
SPECTRA uncovers unconsidered spectral properties. **(a)** Distribution of standard deviation of test metric performance across splits generated with the same spectral parameter for the RIF, PZA, INH, GFP, and SARS-CoV-2 datasets across all tested models. Splits with a high standard deviation could have novel spectral properties. **(b)** The resistance determining region (RRDR) of the *rpoB* gene in Tuberculosis is a 26 amino acid region strongly associated with Rifampicin resistance. **(c)** Diff-RRDR is a metric that measures the difference in the minimum and maximum positions of observed RRDR mutations in the train and test. **(d-e)** The spectral performance curve for CNN **(d)** and ESM2 **(e)** in the RIF dataset (shown left). Test AUROC versus diff-RRDR for CNN **(d)** and ESM2 **(e)** in the RIF dataset (shown right). Highlighted are points representing splits generated at a spectral parameter of 0.90, 0.95, and 1. All test metrics are averaged across three independent runs.

To investigate this hypothesis, we calculated the difference in the range of positions for RRDR mutations observed in the train and the test splits (diff-RRDR) (Figure 4c). Diff-RRDR explains the variance in model performance observed at SP = 0.9 (Spearman’s rank correlation of *ρ* = *−*0.51, p-value=1.79E-5, between diff-RRDR and AUROC, Figure 4d). We observe similar patterns for the SP = 1.0 split and SP = 0.95 split (Figure 4d). This correlation is model specific; ESM2 performance experiences no degradation in performance as diff-RRDR increases in the SP = 0.9, 0.95, 1.0 splits (Figure 4e). ESM2 considers a larger range of input positions (512) than the CNN (12) when making a prediction. We expect this to make ESM2 more invariant to increases in diff-RRDR as has been observed in previous work comparing the performance of LLMs to CNNs and attributing the higher performance of the former to a larger context window around a target token [49]. Our results improve our understanding of *M. tuberculosis* resistance prediction where state-of-the-art performance is currently achieved by CNN models [1]. The generalizability of *M. tuberculosis* resistance prediction models can be improved by using LLMs and potentially other models that consider a longer context of DNA sequence. We characterize the difference in the length of genetic context, e,g, diff-RRDR, as a spectral property relevant for phenotype prediction for proteins where functionally impactful mutations concentrate along nucleotide sequence e.g. in the active site. Hence to fully evaluate generalizability, multiple biologically informed spectral properties should be used when running SPECTRA.

### SPECTRA can help evaluate the generalizability of foundation models in biology

Increasingly machine learning models for molecular sequencing data are pre-trained on large datasets and then trained and tested on usually smaller task-specific datasets not encountered in pre-training [12, 50, 51]. These foundation models have the potential to offer better flexibility and adaptability to a wide range of tasks as they have done in computer vision [52] and natural language processing [53–55]. Despite the potential of these models, their generalizability is unknown as they have rarely been tested prospectively on non-overlapping datasets [56, 57]. Current approaches to benchmark foundation models report the average performance across multiple task-specific datasets. However, this approach exaggerates foundation model generalizability by failing to consider cross-split overlap in the task-specific data splits and the overlap between the pre-training and task-specific datasets. Running SPECTRA with foundation models on task-specific datasets evaluates generalizability by measuring AUSPCs. We can then assess the effect of overlap between the pre-training and task-specific datasets on these AUSPCs to understand foundation model generalizability.

To demonstrate this capability of SPECTRA, we investigate the generalizability of several protein foundation models. From our previous analysis, the AUSPC of ESM2 for RIF, PZA, INH, SARS-CoV-2, and GFP phenotype prediction varied widely from 0.91 in RIF to 0.26 in SARS-CoV-2 (Figure 3b). We calculated the overlap between these task-specific datasets and Uniref50, the pre-training dataset used by ESM2 [58] (Figure 5a, Section 5 Methods) finding significant correlation between this overalp and the AUSPC of ESM2 (Spearman rank correlation = 0.9, P-value = 1.4e-27, Figure 5a). Fine-tuning improves ESM2 AUSPC for PZA, SARS-CoV-2, and GFP phenotypic prediction (Figure 3b). The effect of overlap between pre-training and task-specific data holds when evaluating protein foundation models other than ESM2 including Transception, MSATransformer, ESM1v, and Progen on five molecular sequencing datasets in the ProteinGym benchmark [16] (Supplementary Note 3, Figure 5b; Spearman rank correlation = 0.9, P-value = 0.04).

**Figure 5:**
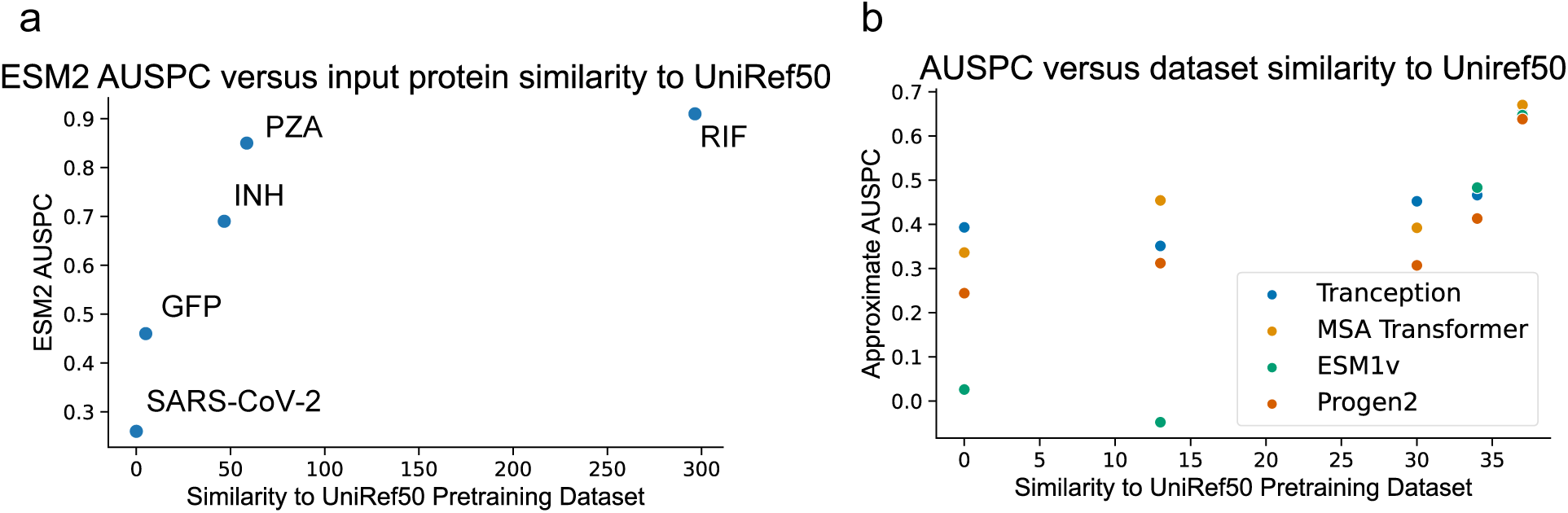
Influence of pre-training set similarity on protein foundation model generalizability. **(a)** ESM2 AUSPC versus similarity to Uniref50, the pretraining dataset of ESM2, for the RIF, INH, PZA, GFP, and Covid tasks. Each point is labeled with the corresponding molecular sequencing dataset. **(b)** Approximate AUSPC versus UniRef50 similarity for four protein foundation models, Transception, MSA Transformer, ESM1v, and Progen2, in five molecular sequencing datsets from ProteinGym. (Dataset names in order from left to right: A4D664-9INFA-Soh-CCL141-2019, A0A140D2T1-ZIKV-Sourisseau-growth-2019, A4-HUMAN-Seuma-2021,AACC1-PSEAI-Dandage-2018, A4GRB6-PSEAI-Chen-2020). All correlations shown are significant (P-value < 0.05).

### Computational cost of SPECTRA

We found running SPECTRA can be computationally expensive depending on (1) the computational complexity of comparing spectral properties, (2) the size of the spectral property graph, and (3) the size of the model to be evaluated. We estimate runtime ranges for generating SPECTRA splits and spectral performance curves with more details in Supplementary Note 4. SPECTRA split generation ranged from 9 hours for PDBBind to 5 minutes for the amyloid beta protein aggregation dataset from ProteinGym. Spectral performance curve generation ranged from 5 days for ESM2-Finetuned to one hour for logistic regression. To improve runtimes we utilized multi-core CPU machines to parallelize spectral property comparison and multiple GPUs to train models on different spectral parameters in parallel (Section 3.2 Methods).

## Discussion

Understanding how well molecular machine learning models perform on unseen data is a fundamental problem for protein design [59, 60], defense against emerging pathogens [34, 61, 62], and therapeutic science [63–65]. Increasingly, studies are demonstrating how reported benchmarking model performance is overly optimistic about model generalizability [66–68]. We show this generalizability gap between model performance on benchmarks and external datasets is due to inadequate assessment of overlap between training, test, and external datasets. We demonstrate that model performance decreases in existing benchmarks as cross-split overlap decreases. To address this challenge, we introduce the spectral framework for evaluating, comparing, and understanding model generalizability by explicitly controlling cross-split overlap when constructing train-test splits to generate spectral performance curves (SPCs) and calculate the area under the spectral performance curve (AUSPC). We demonstrate SPECTRA summarizes model test performance more comprehensively than in previous benchmarks. We show SPECTRA can identify novel and relevant spectral properties or molecular properties that influence model generalizability. We also demonstrate SPECTRA’s ability to be used as a tool to evaluate foundation models in biology. After applying SPECTRA to four protein foundation models across 11 molecular sequencing datasets, we found that foundation models have better generalizability in task-specific datasets with greater similarity to the pre-training dataset. These findings corroborate existing literature, which indicates single-cell foundation models struggle to generalize in task-specific datasets dissimilar from pre-training datasets [56].

Evaluating foundation models in biology across multiple splits is computationally expensive, especially for larger models and datasets. Despite the high computational cost of SPECTRA, SPECTRA has only to be run once to generate splits for a given task. Studies can release the splits generated by SPECTRA as part of new benchmarks to offset the cost of running SPECTRA. Existing SB and MB splits are attractive because they are computationally inexpensive. However, as shown in this paper, the time saved in computation leads to a mischaracterization of model generalizability. Model evaluation should be seen as a step in model development as computationally expensive as model training, not an afterthought to generate a performance metric. The purpose of this study is not to rank models by their AUSPCs but to demonstrate the spectral framework for model evaluation on state-of-the-art models. To rank models requires averaging over many AUSPCs calculated across various tasks in a SPECTRA benchmark for molecular sequencing data.

The choice of the spectral property is pivotal for SPECTRA. This property is ideally informed by the biological mechanism of how the sequence encodes the phenotype or cellular characteristic to be predicted. Further, it is important to confirm that the procedure for capturing whether two samples share a spectral property is robust and biologically meaningful. Assuming so, if the model performance variance is observed to be high within spectral parameter bins, we show that this suggests that the evaluated spectral property is not sufficient for assessing generalizability, and the evaluation of additional spectral properties is advisable. As the community utilizes SPECTRA, we expect standardized definitions of spectral properties will be created for different tasks.

It is important to note we plot model performance against the spectral parameter instead of cross-split overlap because the minimal cross-split overlap (SP = 0) varies across tasks and definitions of spectral properties. The spectral parameter reflects the relative impact on model performance per unit decrease of cross-split overlap, offering a more holistic view of model generalizability. By considering these effects across the entire spectral parameter range, the resulting AUSPC enables comparison of generalizability across different models and tasks.

Our framework makes urgent the need to define and then refine molecular sequence property benchmarks for generalizability. In the case of *M. tuberculosis*’ resistance to rifampicin, previous work has found high AUROC using logistic regression models which supported the development of commercial genotypic assays for rifampicin resistance prediction. SPECTRA found logistic regression has high AUROC at SP = 0 (maximal cross-split overlap), but degrades to near random performance at SP = 1 (no cross-split overlap). This corresponds to current understanding that resistance to rifampicin is encoded by a handful of very common mutations concentrated in the active site and a long tail of more rare mutations in a more distant proximity that can be more common in specific regions of the world. As a result, these rifampicin resistance commercial tests were later recognized to misclassify resistance in large populations of patients in specific geographic regions [69]. SPECTRA could have provided a more realistic understanding of the sensitivity of these tests and the mutations they are based on before deployment. This case of *M. tuberculosis*’ resistance to rifampicin has implications for any protein where mutations with a functional effect on phenotype concentrate along the protein’s length or structure including for example the concentration of immune escape mutations in the SARS-CoV spike receptor binding domain.

SPECTRA can be used with models beyond the molecular sequencing domain. For example, in ML for small molecule therapeutics, the chemical structure is the spectral property used to separate molecules into structurally dissimilar train-test splits [70]. In inverse protein folding problems, when models associate the amino acid sequence that folds into a particular protein structure, the ontological label of protein folds can be used as the spectral property [71]. In patient-level medical datasets, spectral properties can be defined based on patient demographics and clinical characteristics.

In biology, there is a general belief that observations made in the past predict future observations or that models can only generalize to samples that are similar to previous data. Our results challenge this philosophy by demonstrating that in some cases models perform well on sequences with never-before-seen mutations, motifs, or sequence identity. Our work identifies molecular sequence properties that allow models to generalize in these scenarios, such as sequence similarity to a pretraining set. The SPECTRA framework for model evaluation represents a new frontier, generating a robust understanding of model performance beyond a metric on a metadata or similarity-based split.

## Data availability

All data used in this study is publicly available. The data used for the RIF, INH, and PZA datasets can be found in Green *et al.* [1]. The data used for the GFP dataset is from Sarkisyan *et al.* [45]. The data used for the SARS-CoV-2 dataset is from Greaney *et al.* [44]. All other datasets were directly downloaded from their benchmark of origin. All data is also available on the project Github at https://github.com/mims-harvard/SPECTRA and on Harvard Dataverse at https://dataverse.harvard.edu/dataset.xhtml?persistentId=doi:10.7910/DVN/W5UUNN.

## Code availability

The code to reproduce results, together with documentation and examples of usage, are available on GitHub at https://github.com/mims-harvard/SPECTRA.

## Acknowledgements

Y.E. is supported by grant T32 HG002295 from the National Human Genome Research Institute and the NSDEG fellowship. Y.E. and M.Z. gratefully acknowledge the support of NIH R01-HD108794, NSF CAREER 2339524, US DoD FA8702-15-D-0001, awards from Harvard Data Science Initiative, Amazon Faculty Research, Google Research Scholar Program, AstraZeneca Research, Roche Alliance with Distinguished Scientists, Sanofi iDEA-iTECH Award, Pfizer Research, Chan Zuckerberg Initiative, John and Virginia Kaneb Fellowship award at Harvard Medical School, Aligning Science Across Parkinson’s (ASAP) Initiative, Biswas Computational Biology Initiative in partnership with the Milken Institute, and Kempner Institute for the Study of Natural and Artificial Intelligence at Harvard University. Any opinions, findings, conclusions or recommendations expressed in this material are those of the authors and do not necessarily reflect the views of the funders.

## Authors contributions

Y.E. retrieved and processed all data. Y.E. performed all analyses except the spectral performance curve for GearNet and GearNet-finetuned which was performed by A.S. and for ESM2 for the SARS-CoV-2 dataset which was performed by D.B. M.M. assisted with the processing of the Tuberculosis data for the INH, PZA, and RIF datasets. Y.E, M.F., and M.Z. designed the study. All authors contributed to writing the manuscript.

## Online Methods

The Methods describe details for (1) the SPECTRA framework, (2) the curation of all molecular sequencing datasets used in this study, (3) model architecture and training, and (4) characterizing distribution shifts in molecular sequencing datasets.

## 1 The SPECTRA framework for model evaluation

For an input molecular sequencing dataset and model, SPECTRA consists of 3 steps: (1) spectral property definition, (2) spectral property graph construction and split generation, and (3) model spectral performance curve generation.

### 1.1 Defining spectral properties

A molecular sequence property (MSP) is a property that is either given or calculated inherent to a molecular sequence. Spectral properties are MSPs that influence model generalizability. Spectral properties are task and data specific: what may be a spectral property for one data type and task is not one in another data modality and task. For example, for predicting DNA binding motifs, the number of adenines in an input nucleic acid sequence is a molecular sequence property but not a spectral property. For the task of secondary structure prediction, the 3D structure of an amino acid sequence is a MSP which is also a spectral property because the structural motifs present in a train set versus a test set will influence model generalizability.

The choice of how to define spectral properties is dataset and problem-specific. The datasets used in this study can be divided into two categories: (1) mutational scan datasets (MSD), which comprise a single set of sequences with different mutations and their effect on phenotype, and (2) sequence-to-sequence datasets (SD), which comprise of different sequences and their properties.

In SDs, the spectral property of interest is sequence identity. To calculate whether two sequences share sequence identity, we perform a pairwise alignment between input sequences and calculate the proportion of aligned positions to the length of the pairwise alignment. If this proportion is greater than 0.3, then the two sequences share this spectral property [80]. We use Biopython [81] to align with a match score of 1, a mismatch score of -2, and a gap score of -2.5. We employed heuristics to define the comparator for larger datasets when exhaustive pairwise comparison of all sequences was not computationally feasible (Section 5 Methods). In MSDs, phenotypically meaningful differences are in the scale of single mutations. Thus, using the definition of the spectral property from SDs would underestimate differences between samples. To address this, we represent samples in MSDs by their sample barcode or a string representation of the mutations present in the sample. The spectral property of a sample is its sample barcode. Two samples share this property if their sample barcodes share at least one mutation.

### 1.2 Constructing spectral property graphs and SPECTRA split generation

After the spectral property is defined, a spectral property graph (SPG) is constructed where nodes are samples in the input dataset, and edges are between samples that share a spectral property (Supplementary Figure S1-S2). Finding a split such that no two samples share a spectral property is the same as finding the maximal independent set of the SPG or the maximum set of vertices such that no two nodes share an edge [82]. Finding the maximal independent set is NP-Hard [83], we approximate it via a greedy random algorithm where we (1) randomly order SPG vertices, (2) choose the first vertex and remove all neighbors, and (3) continue until no vertices remain in the graph. To create an overlap in generated splits, we introduce the spectral parameter to the algorithm. Instead of deleting every neighbor, we delete each neighbor with a probability equal to the spectral parameter. If the spectral parameter is 1, we approximate the maximal independent set; if it is 0, we perform a random split. Given a set of nodes returned by the independent set algorithm, we produce an 80-20 train-test split. Sample SPGs can be found in Supplementary Figure S1-S2, and statistics for all generated SPGs can be found in Supplementary Table S2.

This procedure is complicated in MSDs where sample barcodes map to multiple samples (i.e., in MSDs where the number of unique sample barcodes is not equal to the number of samples). As a result, if split generation does not consider the number of samples, splits can be generated with a small or uneven distribution of samples (i.e., a train set with 100 samples and a test set with 10000 samples). To address this, we applied two changes: (1) weighing nodes in the SPG by the number of samples represented by the sample barcode and biased the algorithm to choose these nodes, and (2) when splitting the nodes into train-test splits, we ran a subset sum algorithm to ensure train and test splits had 80% and 20% of samples respectively (Supplementary Note 1). Statistics for all generated SPECTRA splits can be found in the supplement (Supplementary Figure S5-S6).

### 1.3 Generating spectral performance curves

To generate a spectral performance curve, we create splits with spectral parameters between 0 and 1 in 0.05 increments. For each spectral parameter, we generate three splits with different random seeds. We then train and test models on generated splits and plot model test performance versus spectral parameters. The area under this curve is the area under the spectral performance curve (AUSPC). We provide spectral performance curves and AUSPCs for all relevant models in the GFP, SARS-CoV-2, RIF, PZA, and INH datasets (Supplementary Figure S7-S11).

## 2 Datasets

This section outlines the datasets and processing performed for this study.

### 2.1 Tuberculosis dataset

Tuberculosis (TB) sequencing and antimicrobial screening data are obtained from Green *et. al* [1]. Paired-end reads are trimmed with trimmomatic [84], assembled using Spades [85] into contigs, and aligned to reference TB reference genome H37Rv via minimap2 [86]. All TB datasets used in this study are MSDs. To generate sample barcodes for input TB isolates, we use Pilon [87] to generate VCF files. Bcftools [88] is then used to pull all variants identified in the regions of interest for a particular drug (Supplementary Table S1). From the output of Bcftools, each variant is summarized as a mutational barcode or a string representation of the position and nucleic acid change that defines the mutation. Each isolate is then summarized with a sample barcode or a concatenation of the mutational barcodes present in the isolate. We collect nucleic acid sequences from the contigs mapped to the regions of interest (Supplementary Note 2). For the Rifampicin resistance prediction task (RIF), we have 17,474 *M. tuberculosis* clinical isolates where 4,963 are resistant, and 12,511 are susceptible. There are 3,998 unique sample barcodes and 2,066 unique mutational barcodes. For Pyrazinamide (PZA), there are 12,146 isolates, where 2,166 are resistant and 9,980 are susceptible. There are 2,571 unique sample barcodes and 2,742 unique mutational barcodes. For Isoniazid (INH), there are 26,574 isolates, where 10,580 are resistant and 15,994 susceptible. There are 4,952 unique sample barcodes and 4,455 unique mutational barcodes.

### 2.2 GFP dataset

GFP dataset is a MPD where amino acid sequences, sample barcodes, and phenotypes are obtained from Sarkisyan *et al.* [45]. The dataset maps amino acid sequences of the green fluorescent protein (GFP) of the *Aequorea victoria* jellyfish to a value representing the fluorescence of the GFP protein. This is a regressive task where fluorescence values are log-transformed, and min-max normalized. GFP has 54,024 samples with 54,024 unique sample barcodes and 1,880 unique mutation barcodes (Supplementary Figure S3). The performance metric is Spearman’s rank correlation between predicted and experimentally measured fluorescence values.

### 2.3 SARS-CoV-2 dataset

SARS-CoV-2 dataset is a MPD where amino acid sequences, sample barcodes, and phenotypes are obtained from Greaney *et al.* [44]. The dataset maps mutations of the amino acid sequences for the receptor binding domain (RBD) of the SARS-CoV-2 spike protein to a value representing vaccine escape. This phenotype is measured by exposing an RBD sequence to a series of human antibodies and reporting the proportion of RBD sequences bound by each antibody. The higher the proportion, the less the mutation in the RBD domain is associated with vaccine escape. We take the smallest bound proportion for each mutated sequence to generate labels and log-transform and min-max normalize the values. This is a regressive task with 438,046 samples with 22,341 unique sample barcodes and 2,391 unique mutation barcodes (Supplementary Figure S4). The performance metric is Spearman’s rank correlation between predicted and groundtruth escape values.

### 2.4 PEER benchmark datasets

PEER [25] is a benchmark consisting of 17 tasks spanning five task categories (protein function prediction, protein localization prediction, protein structure prediction, protein-protein interaction prediction, and protein-ligand interaction prediction). From PEER, we run SPECTRA on the subcellular localization dataset for the protein localization prediction task from Armenteros *et al.* [89], a SD with 13,949 samples which maps protein sequences to one of ten labels that present subcellular locations. This dataset is a classification task reporting per-label/class accuracy as a performance metric.

### 2.5 ProteinGym benchmark datasets

ProteinGym [16] is a benchmark consisting of 94 deep mutational scan datasets assessing the effect of mutations on measured protein properties. All ProteinGym datasets are MPDs. From ProteinGym, we run SPECTRA on the amyloid beta protein aggregation dataset. This dataset from Seuma *et al.* [90] maps mutations of the amyloid beta peptide that aggregates in Alzheimer’s disease to an enrichment score reflecting the ability of the mutated peptide to aggregate in a cell-based selection assay. This is a regressive task with 14,483 sample barcodes. The performance metric is Spearman’s rank correlation between predicted and groundtruth assay readouts. We also run SPECTRA on the RNA recognition motif (RRM) dataset. This dataset from Melamed *et al.* [91] maps mutations in the RRM2 domain of the *Saccharomyces cerevisiae* yeast poly(A)-binding protein (Pab1) to an enrichment score. This score is calculated by taking the proportion of yeast strains with a specific mutation in an input population before and after selection. This is a regressive task with 37,708 sample barcodes. The performance metric is Spearman’s rank correlation between predicted and groundtruth enrichment values.

### 2.6 TAPE benchmark dataset

TAPE [24] is a benchmark consisting of five tasks (secondary structure prediction, contact prediction, remote homology detection, fluorescence landscape prediction, and stability landscape prediction). We run SPECTRA on a dataset from Hou *et al.* [92] for the remote homology detection task, which maps input protein sequences to one of 1,195 different fold classifications. This dataset is a SD with 16,291 samples and is a sequence classification task, reporting average accuracy across labels. We also run SPECTRA on a secondary structure dataset for the secondary structure prediction task from Klausen *et al.* [93], a SD with 11,411 samples, which maps proteins to one of three classes representing different secondary structures. This dataset is a classification task reporting per-label/class accuracy as a performance metric.

### 2.7 PDBBind dataset

The protein data bank bind (PDBBind) dataset [41] is a collection of protein-ligand complexes along with their binding affinities. This is a generative task where models generate a protein-ligand complex from a protein structure and a ligand SMILES structural fingerprint. The performance metric is the root mean square error between the predicted and actual protein-ligand complex [35, 78]. We test two splits of the PDBBind dataset, one from Li *et al.* [67] with 14,993 proteinligand complexes and another from Stärk *et al.* [35] with 16,742 protein-ligand complexes. To run SPECTRA, we first download the dataset from Stärk *et al.* from the provided source and gather protein sequences via the protein data bank (PDB) REST API and use Open Babel [94] to convert ligand Mol2 files to SMI files containing ligand SMILES structural fingerprint. We use the procedure outlined in Li *et al.* to calculate similarity among ligands. We define sequence similarity the same way as the rest of the study (Section 1.1 Methods). We then construct a molecular sequence property graph where every node represents a protein-ligand pair. Edges are between nodes with proteins more than 30% similar or ligands more than 99% similar.

### 2.8 Astex diverse dataset

The Astex diverse dataset [77] is a collection of 85 crystallized protein-ligand pairs used to benchmark binding models. We download the Astex diverse set from the Cambridge Crystallographic Data Centre [95]. To extract ligand smile structures, we use Open Babel to convert ligand Mol2 files to SMI files containing ligand smile structures. To extract protein sequences, we create a custom Python parser to pull protein sequences from protein Mol2 files.

### 2.9 Posebusters dataset

The Posebusters dataset [66] is a collection of 428 protein-ligand pairs used to benchmark binding models. It was created to uncover model performance when tested on dissimilar protein-ligand pairs according to PDBBind. We download the Posebusters dataset, gather protein sequences via the protein data bank (PDB) REST API, and use Open Babel [94] to convert ligand Mol2 files to SMI files containing ligand SMILES structural fingerprints.

## 3 Training models

This section outlines the inputs, architecture, and training details of the machine learning models used in this study.

### 3.1 Model architectures and inputs

#### Logistic Regression

Logistic regression architecture and training is based on Chen *et al.* [96]. This model utilizes one-hot encoded vectors to represent samples, where vector positions indicate the presence of a specific mutational barcode found in training samples. A logistic regression model then fits onto one-hot encoded vectors to predict the interest phenotype.

#### CNN

CNN architecture and training is based on Green *et al.* [1]. This model utilizes one-hot encoded vectors to represent nucleic acid and amino acid sequences where vector positions indicate the presence or absence of a base pair or amino acid. The architecture is modified to take in unaligned sequences where sequences are padded to the length of the longest input sequence.

#### ESM2

ESM2 pretrained model is from Lin *et al.* [12] (650 million parameter version) and is used to generate protein embeddings for input sequences. We chunk up input sequences longer than 512 amino acids, embed each chunk, and average the embeddings. If the dataset has multiple protein sequences as input, we embed each input protein sequence and average before prediction. We convert nucleic acid sequences to protein. We tokenize sequences by amino acid identity before input into ESM2. Once input sequence embeddings are obtained, we train a linear probe [97], a logistic regression model, on input embeddings to predict phenotype. To finetune ESM2, we freeze the first 30 layers of ESM2 and replace the masked language head with a linear layer to predict phenotype. We then train the modified ESM2 to predict phenotype.

#### EVE

EVE architecture is from Frazer *et al.* [33].To construct the MSAs necessary for EVE, we use Jackhmmr [98] to pull sequences from UniRep100 [58] similar to the wildtype sequence for the GFP protein and resistance binding domain of the SARS-CoV-2 spike protein. We then use Muscle to align pulled sequences and the codebase from EVCouplings [99] and EVE [33] to process and filter MSAs. We then train EVE with default suggested parameters. The input for EVE is the input sequence aligned with pulled sequences from UniRep100. EVE then returns a low-dimensional representation of the input MSA, which is then used to predict phenotype via a linear probe [97].

#### SeqDesign

SeqDesign architecture is from Shin *et al.* [46]. Seqdesign input processing is the same as EVE except as input, it takes in raw unaligned sequences of the input sequence with pulled sequences from UniRep100.

#### GearNet training

GearNet architecture is from Zhang *et al.* [8]. The GearNet model is a graph neural network that learns protein representations from the 3D structure of the protein. We utilize a pre-trained GearNet model to generate embeddings using protein structures. Structures are generated from protein sequences using ESMFold [12]. The structures are then passed into the pre-trained GearNet model to create embeddings of size 512. We generate output predictions from the graph-level embeddings by training a linear probe [97] to predict the phenotype of interest. A finetuned GearNet model is also trained on each dataset. Both pre-trained and finetuned Gearnet models are trained and evaluated on the GFP and the SARS-CoV-2 datasets.

### 3.2 Training details

We use suggested hyperparameters from source studies to train all models except otherwise noted. Our objective function for models trained on the GFP and SARS-CoV-2 datasets is mean absolute error; for the RIF, INH, and PZA datasets, it is binary cross-entropy. All models were trained on 1 Tesla A10 except ESM2-Finetuned, which was trained on 4 TeslaA100s on an azure cluster. When applicable, we leverage Weights & Biases [100] to select optimal hyperparameters via a random search for each model over learning rate. All code is written in PyTorch [101].

## 4 Uncovering spectral properties in molecular sequencing datasets

Ultimately, choosing a spectral property should capture domain-specific knowledge about the MSPs learned by models during training. However, SPECTRA can detect whether there exists an unconsidered spectral property. This occurs if large variations exist in model performance in splits generated with the same spectral parameter or if there is a positive slope in the shape of the spectral performance curve (i.e., model performance improves with decreasing cross-split overlap). In our study, we focus on diff-RRDR to explain the variance observed in the spectral performance curve of the CNN in the rifampicin resistance prediction task in *M. tuberculosis*. To calculate diff-RRDR for a train-test split, we identify all positions in each split where a mutation occurred in the resistance-determining region (RRDR) of the RNA polymerase beta-subunit (rpoB) gene. diff-RRDR is determined by finding the difference between the maximum position observed in the train set compared to the test set and likewise for the minimum positions, then adding these differences together, as shown below:

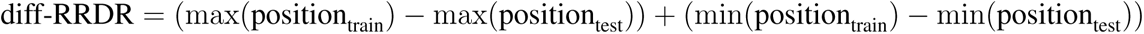

## 5 Using SPECTRA to evaluate foundation models in biology

Beyond the cross-split overlap in evaluation datasets, the cross-split overlap between pre-training and evaluation datasets influences model performance for foundation models. The protein foundation model we evaluated with SPECTRA, ESM2, is pre-trained with Uniref50 [58] with over 60 million clusters of sequences. Each cluster in Uniref50 has a representative sequence that is at least 50% similar to all cluster sequences. To understand what level of similarity between two sequences is significant in Uniref50, we sample one hundred thousand random pairs of representative sequences and calculate the distribution of average random pairwise similarity (Supplementary Figure S12). Two sequences are similar if the sequence similarity, or the proportion of aligned positions in a pairwise alignment, is greater than two standard deviations above mean random pairwise similarity or a sequence similarity of 0.4.

Calculating the sequence similarity between a sequence of interest and UniRef50 representative sequences is computationally infeasible. However, most representative sequences will not be similar to an input sequence. By finding clusters with annotations similar to the protein encoded by the input sequence, we can select the clusters most similar to the input sequences. Once similar clusters are identified, we calculate sequence similarity between the input sequence and the representative sequences of selected clusters and count the number of clusters with sequence similarity greater than 0.4. The number of similar clusters represents the similarity of the input sequence to UniRef50. For tasks with multiple input proteins, we average this number across sequences. The names and similarities calculated for all sequences in this study can be found in Supplementary Table S3 and S4.

## Footnotes

1 The use of the word spectral here refers only to the framework for model evaluation and should not be confused with other uses of the term in matrix analysis.

