## Supplemental Materials for "Evaluating generalizability of artificial intelligence models for molecular datasets"

This PDF file includes:

- Supplementary Notes
- Supplementary Figures
- Supplementary Tables

**Supplementary Note 1: Subset sum algorithm to balance sample numbers in train-test splits.**

Mutational scan datasets (MSDs) comprise of a single set of sequences with different mutations and their effect on phenotype. Samples in MSDs are represented by their sample barcode or a string representation of the mutations present in the sample. In MSDs, multiple samples can share sample barcodes. This occurs when multiple samples have the same mutations. SPECTRA will partition sample barcodes into train and test sets. However, in doing so the number of samples in train and test sets can be skewed. We want to identify a subset of sample barcodes that ensures the intended (80:20) proportion of sample barcodes and number of samples. This is an instance of the subset sum problem which seeks to identify if within a set of numbers a subset exists which sums to a target number [1]. Since this is a NP-Hard problem [2], we approximate the subset sum (Algorithm S1). In words when given a sample barcode, we include it in the training set with a probability of $1-\gamma$, provided that adding the barcode won't surpass the expected proportion of $1-\gamma$ for the total number of samples in the training set. Likewise for the test, except with probability and proportion of $\gamma$.

**Algorithm 1:** SubsetSum approximation algorithm to balance generated SPECTRA train-test splits

**Input:** Selected sample barcodes from spectral property graph $S$, number of samples per barcode $N$, proportion of sample barcodes in the test set $\gamma$, random number generator $r$

**Output:** Balanced Train and Test Sets

$n_{tr} = (1 - \gamma) \sum_{\forall n \in N} n$;
$n_{te} = \gamma \sum_{\forall n \in N} n$;
$sum_{tr} = 0$;
$sum_{te} = 0$;
Initialize the sets train, test, and discarded as empty sets;
**for** $s_i$ *in* $S$ **do**
    $p = r()$;
    **if** $p < 1 - \gamma$ **then**
        **if** $n_{s_i} + sum_{tr} < n_{tr}$ **then**
            Add $s_i$ to train;
            $sum_{tr} += n_{s_i}$;
        **else**
            Add $s_i$ to discarded;
        **end**
    **else**
        **if** $n_{s_i} + sum_{te} < n_{te}$ **then**
            Add $s_i$ to test;
            $sum_{te} += n_{s_i}$;
        **else**
            Add $s_i$ to discarded;
        **end**
    **end**
**end**
**for** $s_i$ *in discarded* **do**
    $p = r()$;
    **if** $p < 1 - \gamma$ **then**
        Add $s_i$ to train;
        $sum_{tr} += n_{s_i}$;
    **else**
        Add $s_i$ to test;
        $sum_{te} += n_{s_i}$;
    **end**
**end**
**return** train, test;

**Supplementary Note 2: Algorithm to pull sequence of interest from *M. tuberculosis* align-**

**ments.**

The coordinates for a gene in the reference *M. tuberculosis* genome, H37Rv, may not correspond to those of a *M. tuberculosis* sample due to mutations before and after the gene that can modify gene coordinates. To address this and collect nucleic acid sequences of regions of interest interest, we process the BAM file that resulted from the pairwise alignment of the input *M. tuberculosis* sample to H37Rv via pysam, a wrapper of samtools [3]. Pysam is used to find the number of contigs associated with a region. If there is one contig, we then search the contig for the first and last 20 reference basepairs of the region of interest. If a match is found, we pull everything in between as the sequence of interest. If we can not find the first and last 20 basepairs (in case of a mutation in these regions), we then repeat the process with basepairs 20-40 from the H37Rv reference. We continue this 20 basepair progression until the sequence is found.

**Supplementary Note 3: Approximating AUSPCs.**

Characterizing SPCs and calculating AUSPCs requires computational resources and time. We identify two instances where AUSPCs can be approximated. Let $f_i(s)$ describe the test performance of model $i$ at SPECTRA splits generated at a spectral parameter of $s$ (SP $= s$). Then the AUSPC of model $i$ is equal to integral of the SPC over all spectral parameters or $\int_0^1 f_i(s)\,ds$.

In the first instance where all samples in MSDs contain a single unique mutation and the spectral property of interest is shared mutations, the spectral performance curve will be a horizontal line. This is because regardless of separation parameter, SPECTRA will always generate splits with no cross-split overlap. As a result the AUSPC reduces: $AUSPC = \int_0^1 f_i(s)\,ds = f_s * (1-0) = f_s$ where $s$ is any spectral parameter. In these special cases AUSPC is equal to test performance on any split. This is the case for the 4D664-9INFA-Soh-CCL141-2019, A0A140D2T1-ZIKV-Sourisseau-growth-2019, A4-HUMAN-Seuma-2021,AACC1- PSEAI-Dandage-2018, and A4GRB6-PSEAI-Chen-2020 datasets from ProteinGym [4].

In the second instance when the objective is to compare model AUSPCs in a benchmark, it is not necessary to characterize the full SPC. The SPC is a monotonically decreasing function or a function that will always decrease or stay constant. As a result the SPC for any model $i$ will be bounded by the test performance at splits generated by a spectral parameter of 0 and 1 or $f_i(1) \leq \int_0^1 f_i(s)\,ds \leq f_i(0)$. With this in mind when comparing two models $i$ and $j$ if $f_i(0) > f_j(0)$ and $f_i(1) > f_j(1)$ then the AUSPC of model $i$ must be greater than that of model $j$. If $f_i(0) = f_j(0)$ and $f_i(1) > f_j(1)$ or $f_i(0) > f_j(0)$ and $f_i(1) = f_j(1)$ then the AUSPC of model $i$ must be greater than that of model $j$. If $f_i(0) > f_j(0)$ and $f_i(1) < f_j(1)$ or vice-versa nothing can be concluded about the relation between AUSPC of model $i$ and $j$, a full SPC must be generated.

This approximation decreases the computational burden when using SPECTRA to calculate AUSPC to compare models. However, this can only approximate the relationship between the AUSPC of two models, they cannot be used to uncover unconsidered spectral properties.

**Supplementary Note 4: Factors that influence runtime of SPECTRA.**

SPECTRA computational cost scales with (1) the computational complexity of comparing spectral properties, (2) the size of the spectral property graph (SPG), and (3) the size of the model to be evaluated.

For the computational complexity of comparing spectral properties, SPECTRA exhibits faster performance with mutation scan datasets (MSDs) compared to sequence to sequence (SDs) due to the computational efficiency of comparing mutational barcodes versus constructing pairwise alignments for spectral property comparison. This is why constructing SPECTRA splits for GFP, a MSD with 54,024 samples, takes 3 hours while construction for PDBBind, a SD with 18,778 samples, takes 9 hours.

The size of the SPG also plays a role. Within MSDs, SPECTRA was able to generate splits faster for PZA (2,571 nodes and 936,431 edges in SPG) than RIF (3,998 nodes and 2,959,746 edges in SPG) on a single-core CPU machine. Within SDs, SPECTRA takes longest for PDBBind (18,778 nodes, 1,987,949 edges in SPG), taking 9 hours to generate 40 splits on a 20-core CPU machine.

After splits are generated, SPECTRA runtime is dictated by the size of the model to be evaluated. To produce the spectral performance curve for the GFP dataset for the largest model considered, ESM2-Finetuned, SPECTRA takes five days for 20 splits across four GPUs. The spectral performance curve for logistic regression for the same dataset takes one hour for 20 splits on a single GPU.

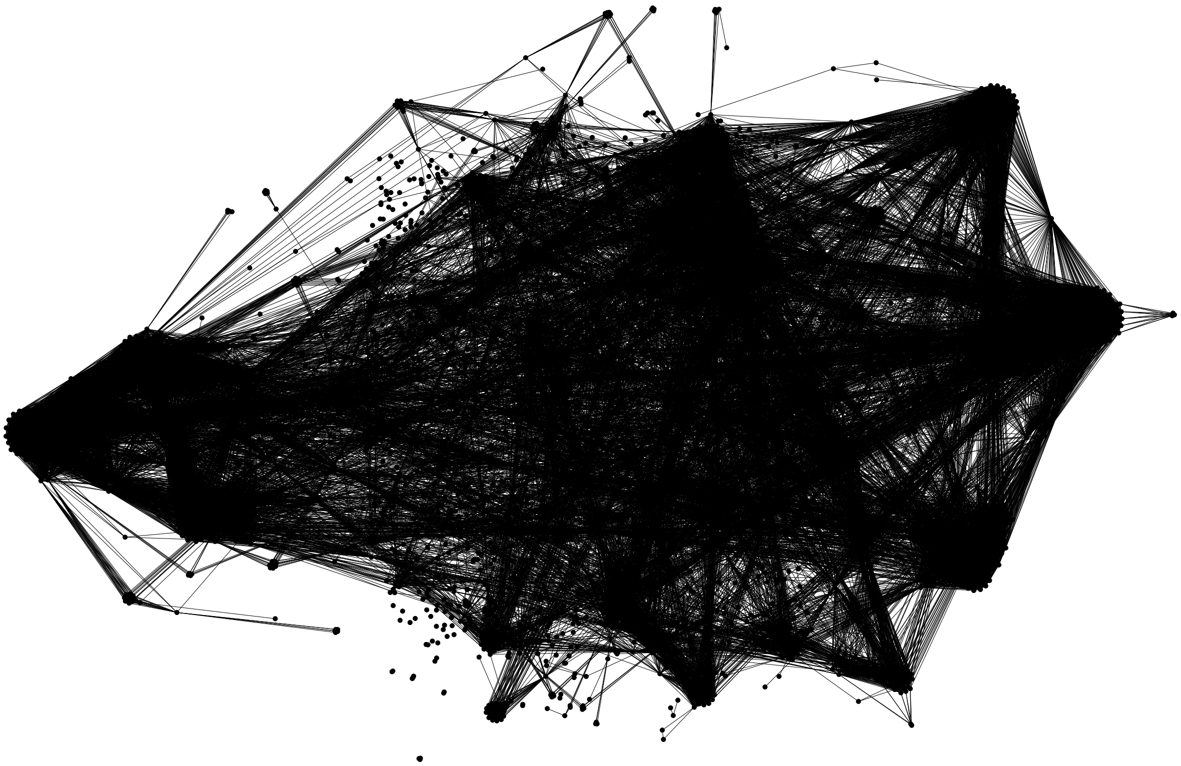

**Figure S1: RIF spectral property graph** Spectral property graph for RIF. Nodes are *M. tuberculosis* samples and edges are between samples that share at least one mutation. There are 3,998 nodes (one for each unique set of mutations) and 2,959,746 edges. The densely connected graph structure reflects how RIF resistance is defined by a small set of key mutations [5].

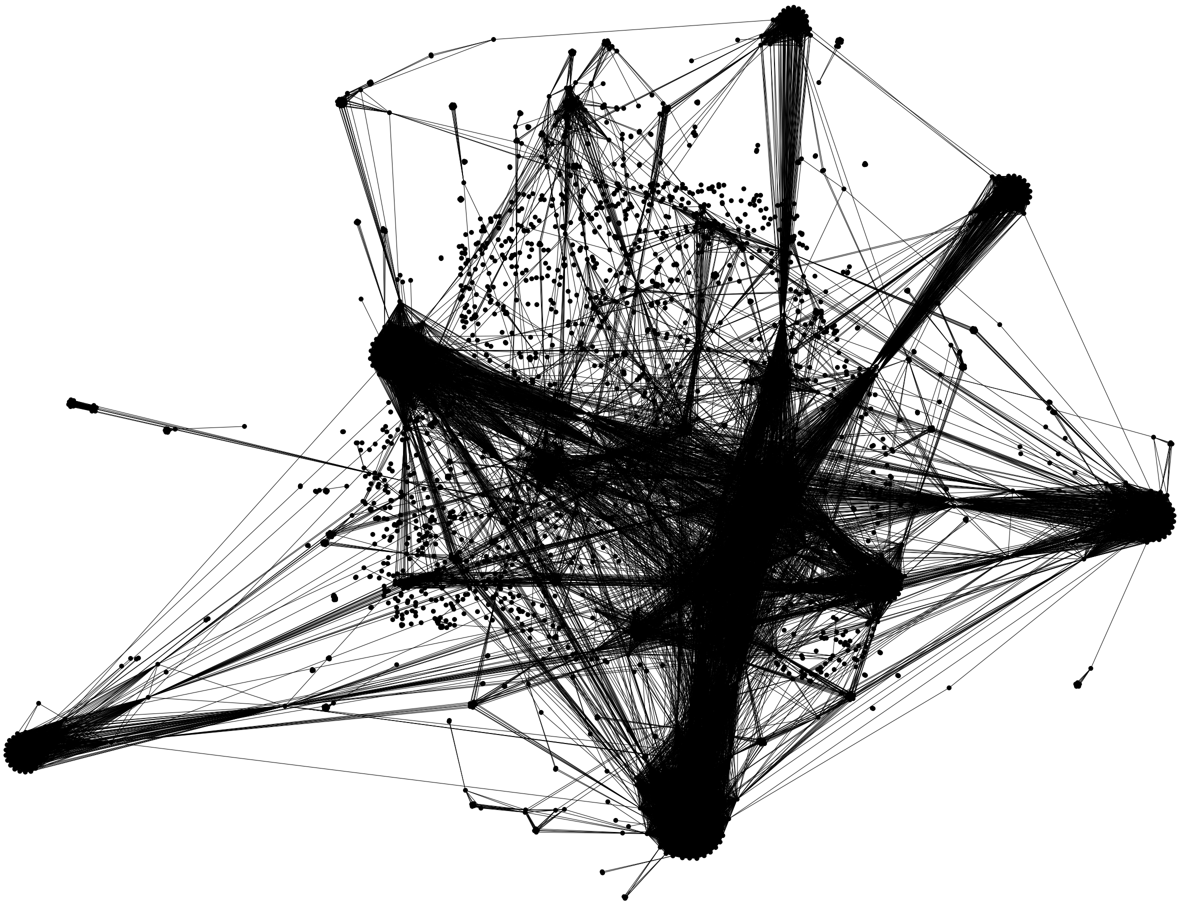

**Figure S2: PZA spectral property graph** Spectral property graph for PZA. Nodes are *M. tuberculosis* samples and edges are between samples that share at least one mutation. There are 2,571 nodes (one for each unique set of mutations) and 936,431 edges. The sparsely connected graph structure reflects how PZA resistance is defined by less-common mutations [6].

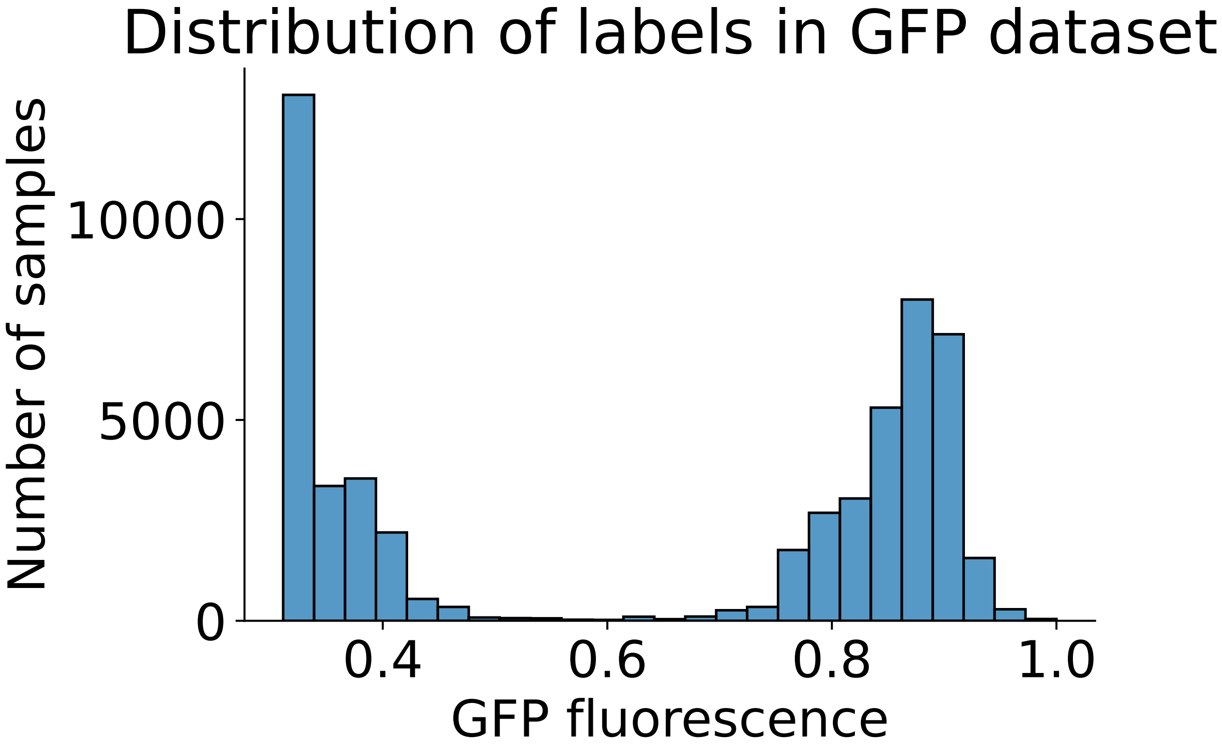


**Figure S3: Distribution of GFP Labels** Distribution of labels for the GFP dataset. The distribution is bimodal where in one one end mutations cause little fluorescence in the GFP while on the other end mutations cause an increase in fluorescence. This distribution was log and min-max normalized to assist in model training.

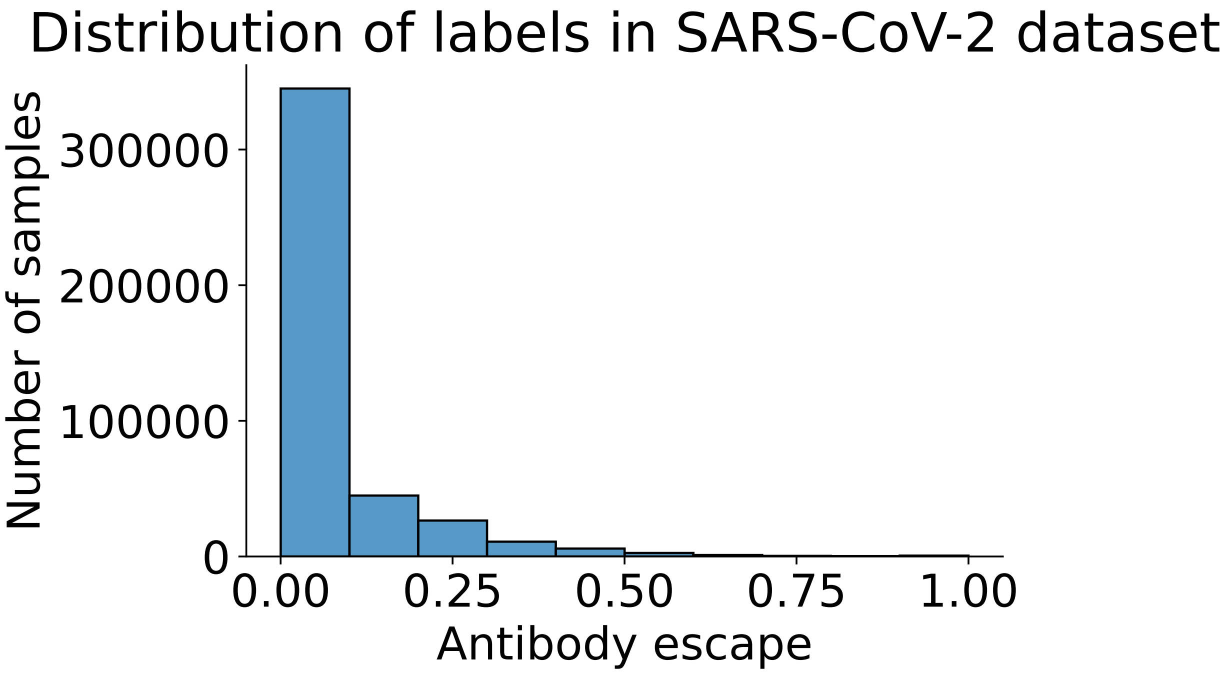


**Figure S4: Distribution of SARS-CoV-2 Labels** Distribution of labels for the SARS-CoV-2 dataset. The distribution is skewed right where the majority of mutations cause little to no antibody escape. This distribution was log and min-max normalized before training to ensure models do not consistently predict no antibody escape due to the skew in the label distribution.

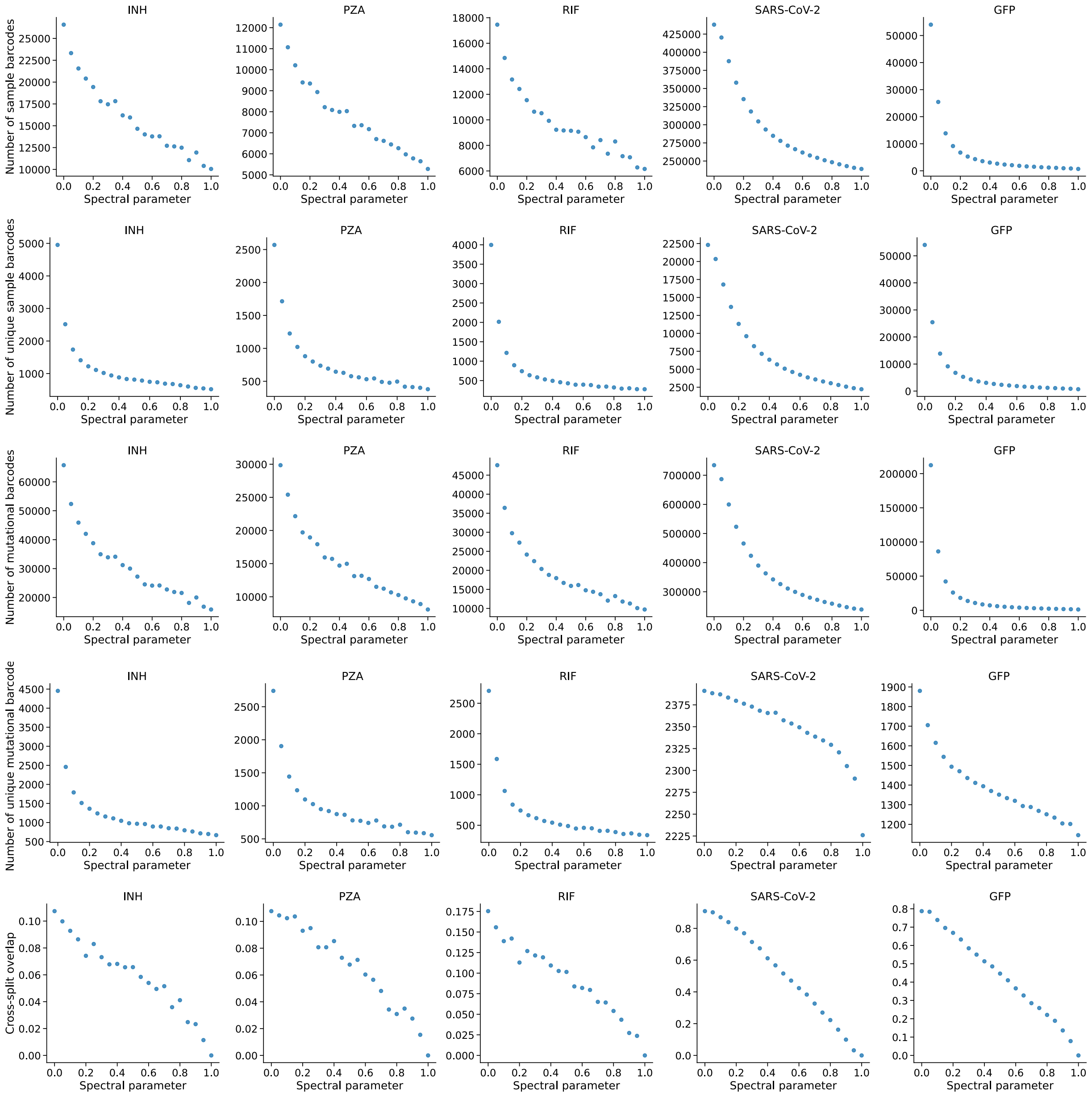


**Figure S5: Statistics for SPECTRA splits generated for INH, PZA, RIF, SARS-CoV-2, and GFP datasets.** Number of sample barcodes, unique sample barcodes, mutational barcodes, unique mutational barcodes, and cross-split overlap for SPECTRA splits. All values shown are averaged across three runs of SPECTRA with different random seeds. Across all datasets, cross-split overlap decreases as spectral parameter increases. The number of samples also decreases, meaning dataset sizes become much smaller at high spectral parameter. In mutation sequence datasets (MSDs) there is a discrepancy between the number of sample barcodes and the unique number of sample barcodes, implying increased dataset size will not always contribute to the diversity of observed mutations.

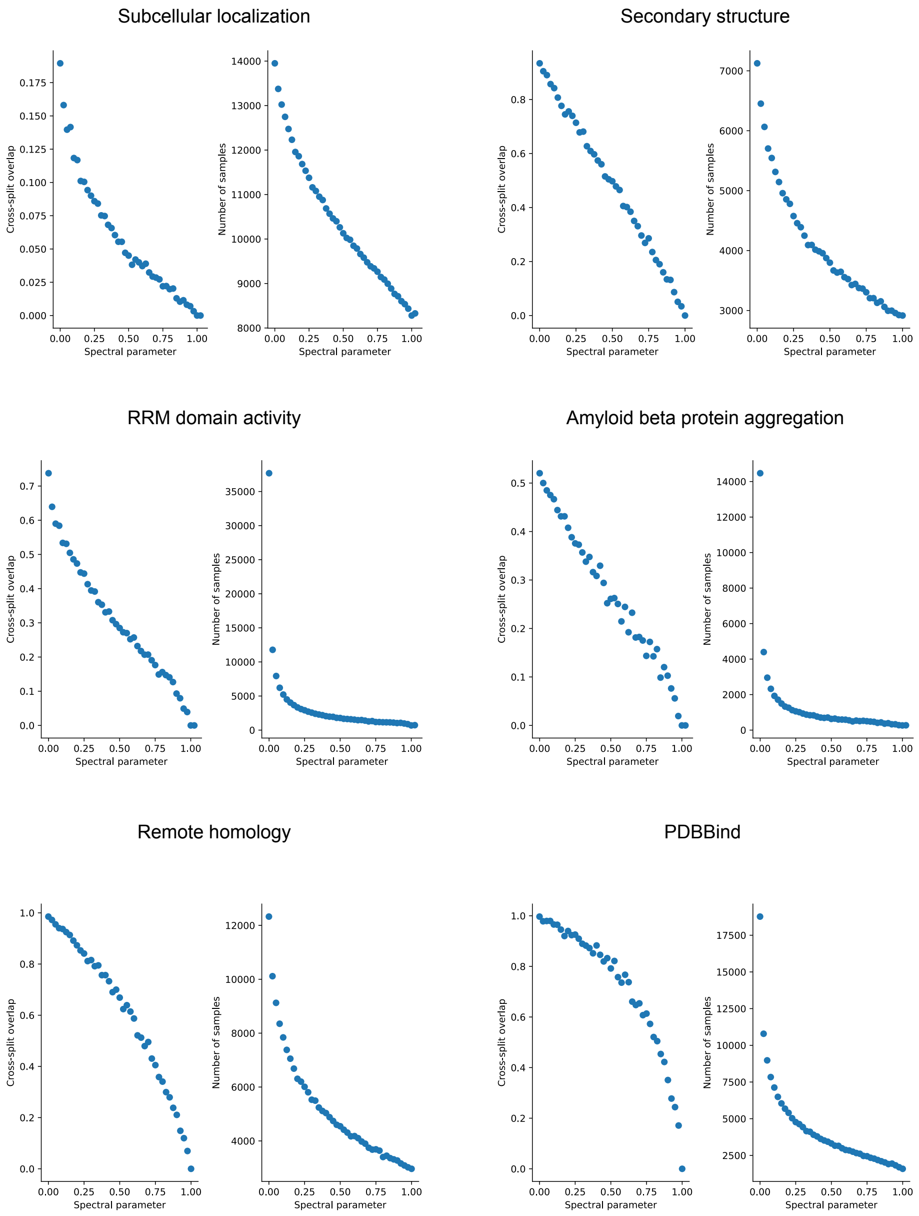


**Figure S6: Statistics for SPECTRA splits generated for PEER subcellular localization, ProteinGym amyloid beta protein aggregation, ProteinGym RRM domain activity, TAPE secondary structure, TAPE remote homology, and PDBBind datasets.**

Cross-split overlap and number of samples for SPECTRA splits for datasets from PEER, TAPE, ProteinGym, and PDBBind. Across all datasets cross-split overlap and sample number decreases as spectral parameter increases. However, the shape of this decrease differs dataset to dataset. This reflects the different structure of the underlying spectral property graph (SPG). In more densely connected SPGs, decreases in number of samples will be more pronounced since more nodes will be deleted at every step of SPECTRA. Also the amount of cross-split overlap at a spectral parameter of zero differs dataset to dataset. This motivates the use of a spectral parameter as it captures the effect on model performance with respect to different amounts of cross-split overlap at random splits.

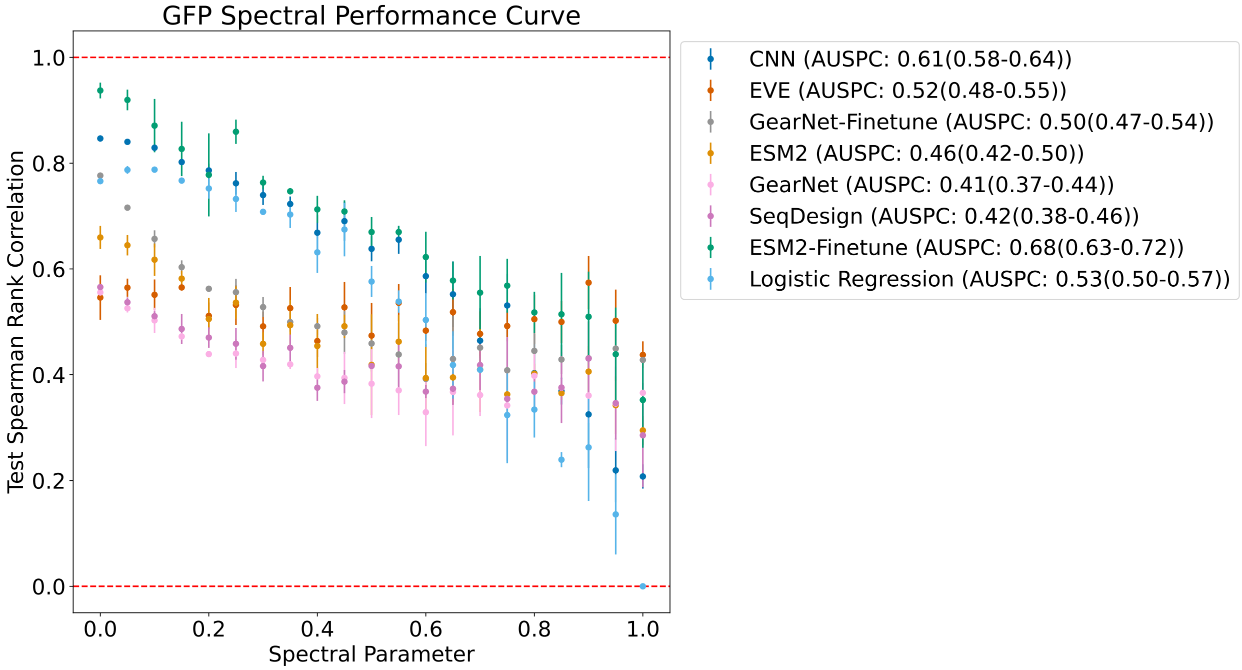


**Figure S7: Spectral performance curve for all models in GFP dataset along with respective AUSPC with confidence interval.**

Points represent the average model test performance at SPECTRA splits generated at the corresponding spectral parameter. Three splits were generated for every value of the spectral parameter. Models were trained three times for every split generated by SPECTRA. Error bars represent the variance in model test performance across the three generated SPECTRA splits. Relatively large error bars indicate the potential presense of unconsidered spectral properties.

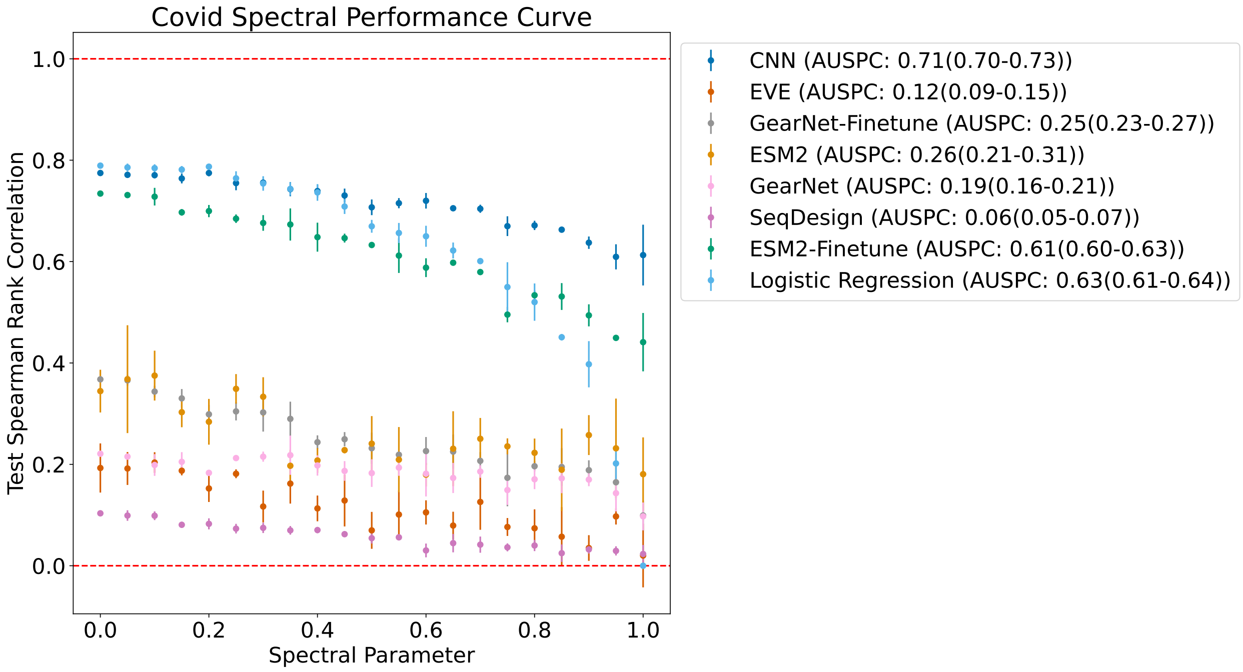


**Figure S8: Spectral performance curve for all models in SARS-CoV-2 dataset along with respective AUSPC with confidence interval.**

Points represent the average model test performance at SPECTRA splits generated at the corresponding spectral parameter. Three splits were generated for every value of the spectral parameter. Models were trained three times for every split generated by SPECTRA. Error bars represent the variance in model test performance across the three generated SPECTRA splits. Relatively large error bars indicate the potential presense of unconsidered spectral properties.

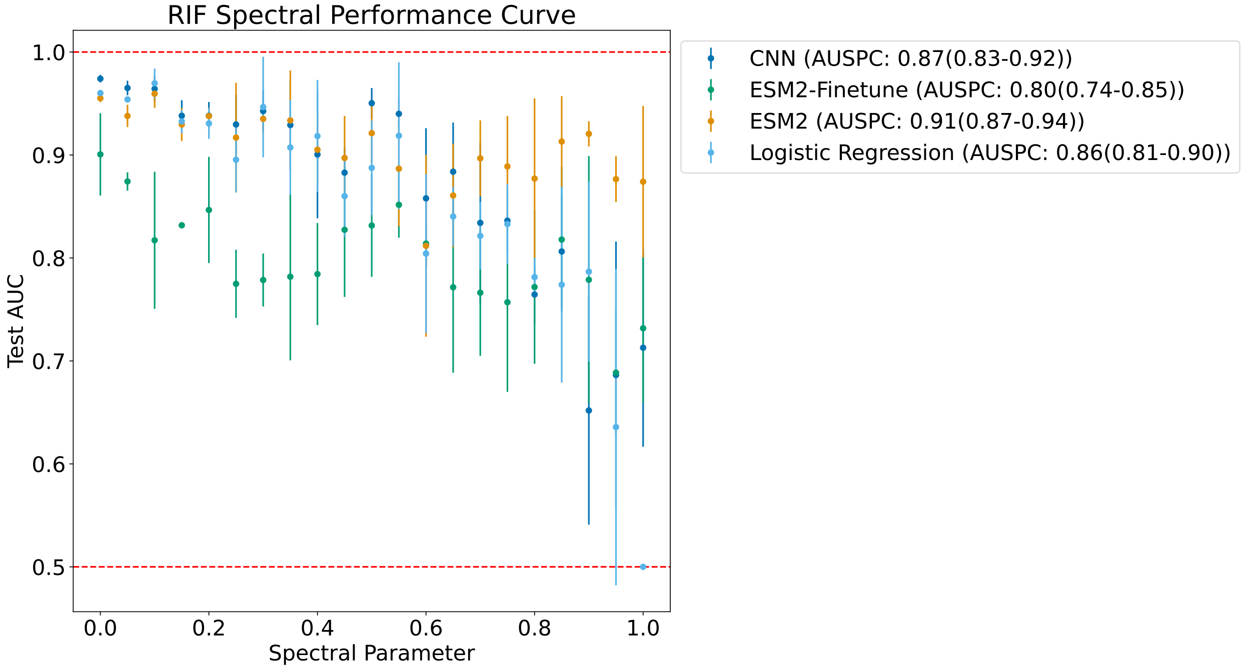


**Figure S9: Spectral performance curve for all models in RIF dataset along with respective AUSPC with confidence interval.**

Points represent the average model test performance at SPECTRA splits generated at the corresponding spectral parameter. Three splits were generated for every value of the spectral parameter. Models were trained three times for every split generated by SPECTRA. Error bars represent the variance in model test performance across the three generated SPECTRA splits. Relatively large error bars indicate the potential presense of unconsidered spectral properties.

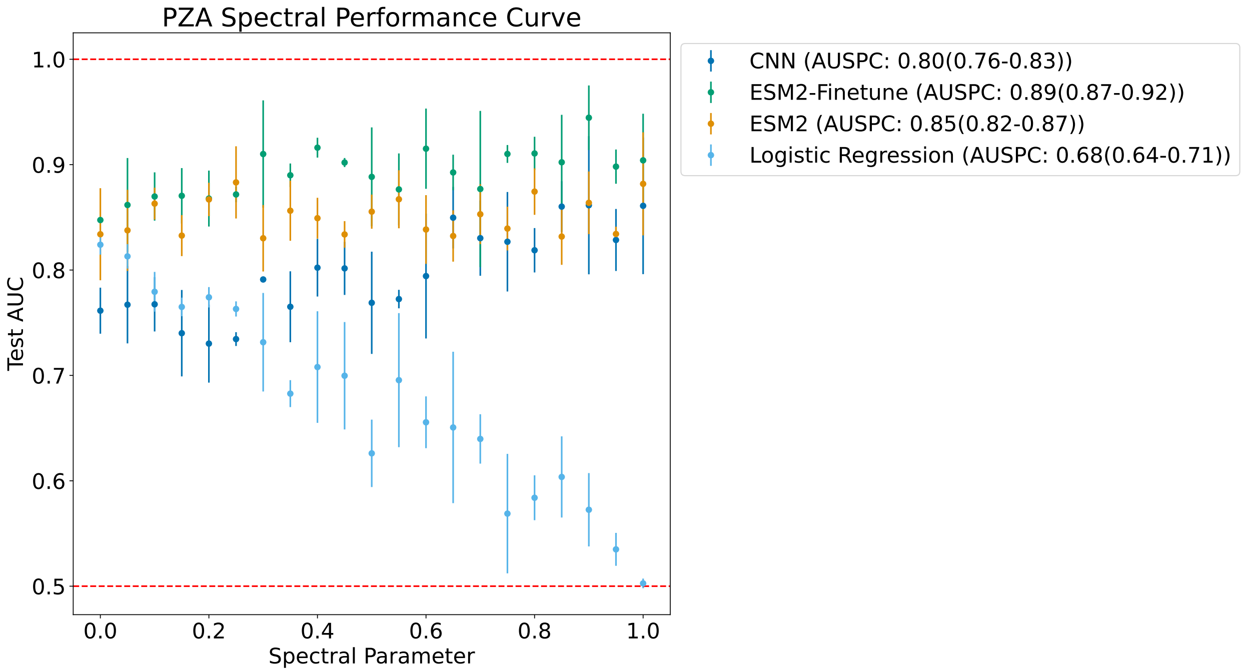


**Figure S10: Spectral performance curve for all models in PZA dataset along with respective AUSPC with confidence interval.**

Points represent the average model test performance at SPECTRA splits generated at the corresponding spectral parameter. Three splits were generated for every value of the spectral parameter. Models were trained three times for every split generated by SPECTRA. Error bars represent the variance in model test performance across the three generated SPECTRA splits. Relatively large error bars indicate the potential presense of unconsidered spectral properties.

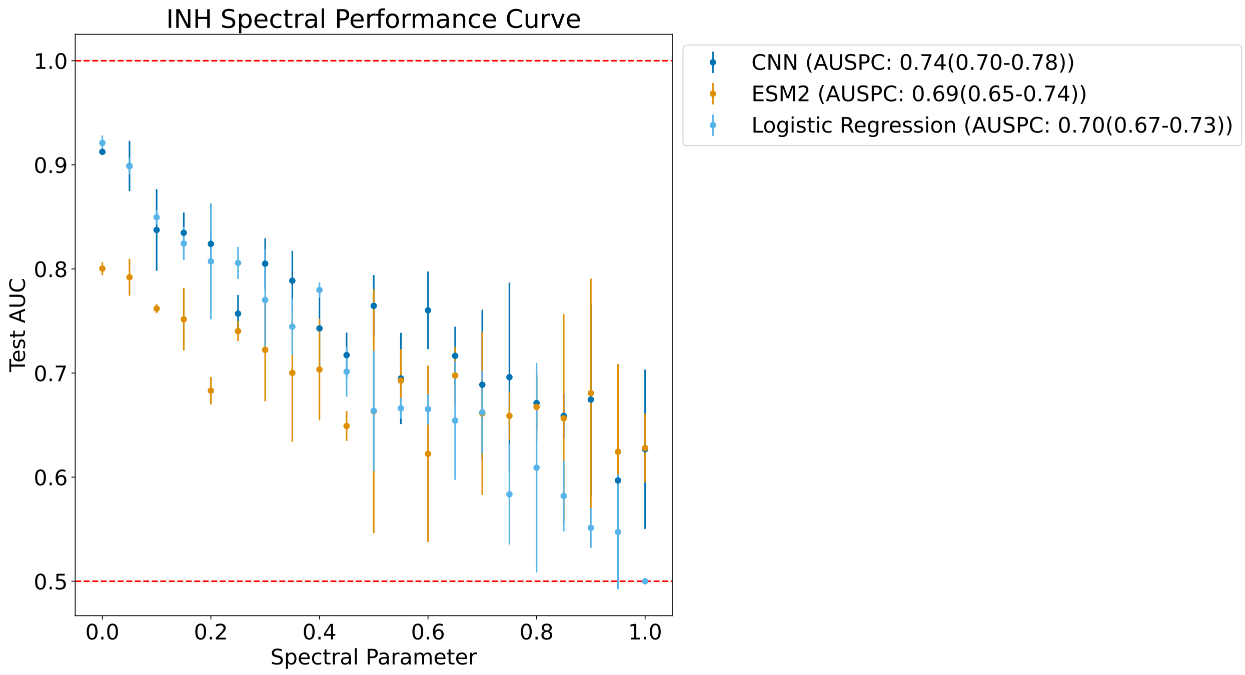


**Figure S11: Spectral performance curve for all models in INH dataset along with respective AUSPC with confidence interval.**

Points represent the average model test performance at SPECTRA splits generated at the corresponding spectral parameter. Three splits were generated for every value of the spectral parameter. Models were trained three times for every split generated by SPECTRA. Error bars represent the variance in model test performance across the three generated SPECTRA splits. Relatively large error bars indicate the potential presense of unconsidered spectral properties.

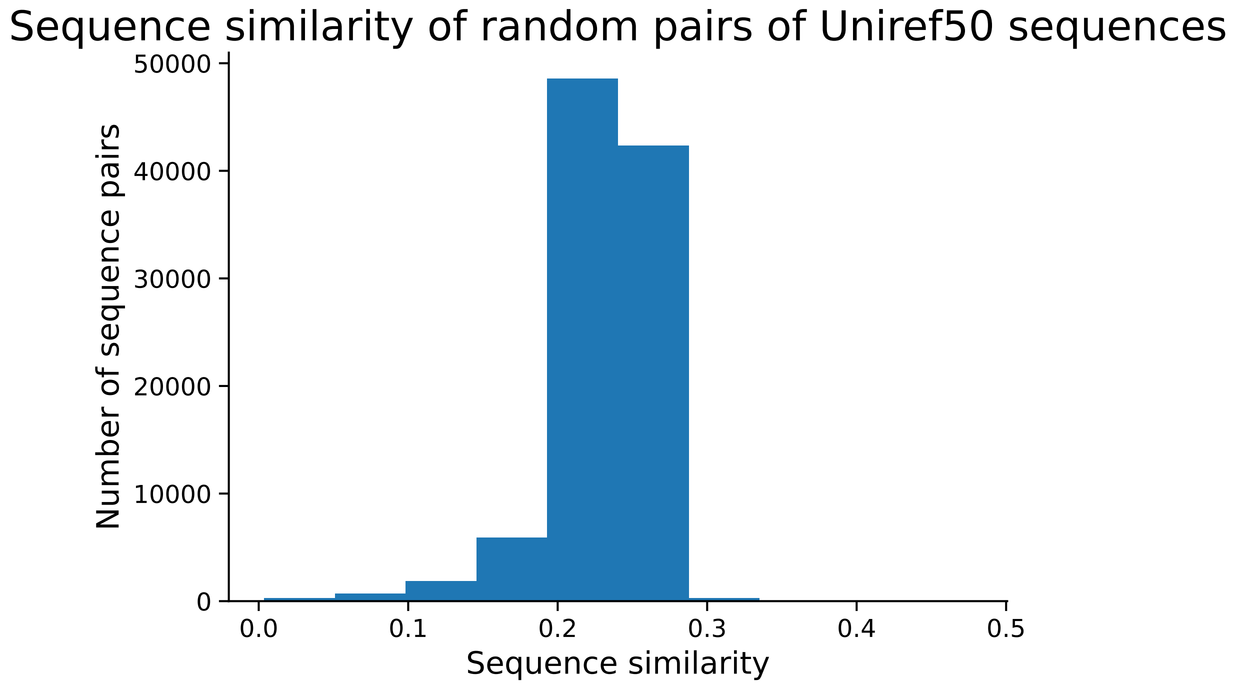


**Figure S12: Sequence similarity of 100000 pairs of sequences from UniRef50.**
Sequence similarity was calculated for 100000 random pairs of cluster representative sequences from UniRef50. A threshold of 0.4, representing two standard deviations away from the mean, was used to denote similarity among cluster representative sequences.

| Drug | Name | Start Coordinate | End Coordinate |
|---|---|---|---|
| PZA | pncA | 2288681 | 2289241 |
| PZA | pncA noncoding | 2289242 | 2289301 |
| PZA | clpC1 | 4038158 | 4040704 |
| PZA | clpC1 noncoding | 4040705 | 4040819 |
| PZA | panD | 4043862 | 4044281 |
| PZA | panD noncoding | 4046119 | 4046239 |
| PZA | Rv1258c | 1406081 | 1407340 |
| PZA | Rv1258c noncoding | 1407341 | 1407407 |
| PZA | PPE35 | 2167649 | 2170612 |
| PZA | PPE35 noncoding | 2170613 | 2170743 |
| PZA | Rv3236c | 3611959 | 3613116 |
| PZA | Rv3236c noncoding | 3613117 | 3613663 |
| RIF | rpoA | 3877464 | 3878507 |
| RIF | rpoA noncoding | 3878508 | 3879052 |
| RIF | rpoB-rpoC | 759807 | 767320 |
| RIF | rpoB-rpoC noncoding | 759420 | 759806 |
| RIF | Rv2752c | 3064515 | 3066191 |
| RIF | Rv2752c noncoding | 3066192 | 3066221 |
| INH | mshA | 575348 | 576790 |
| INH | mshA noncoding | 574670 | 575347 |
| INH | Rv1258c | 1406081 | 1407340 |
| INH | Rv1258c noncoding | 1407341 | 1407407 |
| INH | fabG1-katG | 1673440 | 1675011 |
| INH | fabG1-katG noncoding | 1673336 | 1673439 |
| INH | furA-katG | 2153889 | 2156592 |
| INH | furA-inhA noncoding | 2156593 | 2156652 |
| INH | ndh | 2101651 | 2103042 |
| INH | ndh noncoding | 2103043 | 2103147 |
| INH | ahpC | 2726193 | 2726780 |
| INH | ahpC noncoding | 2726053 | 2726192 |
| INH | Rv2752c | 3064515 | 3066191 |
| INH | Rv2752c noncoding | 3066192 | 3066221 |

**Table S1:** Name, start and end coordinates for genomic regions previously found to be relevant to Pyrazinamide (PZA), Rifampicin (RIF), and Isoniazid (INH) antibiotic resistance in *M. tuberculosis*. Sequences from these regions were pulled as input for all *M. tuberculosis* antibiotic resistance models used in this study. All coordinates are relative to *M. tuberculosis* H37Rv reference strain. [7]

| Dataset | Number of nodes | Number of edges | Number of connected components |
|---|---|---|---|
| Remote homology | 12355 | 223899 | 350 |
| Secondary structure | 7203 | 43349 | 228 |
| Subcellular localization | 13949 | 55130 | 7557 |
| RRM domain activity | 37708 | 5269784 | 134 |
| Amyloid beta protein aggregation | 14483 | 2139791 | 216 |
| RIF | 3998 | 2959746 | 160 |
| PZA | 2571 | 936431 | 294 |
| INH | 4952 | 4587993 | 376 |
| SARS-CoV-2 | 22341 | 919751 | 11 |
| GFP | 54024 | 42134887 | 8 |
| PDBBind | 18778 | 1987949 | 247 |

**Table S2:** Spectral property graph statistics (number of nodes, edges, and connected components), for spectral property graphs generated in this study.

| **Gene (Associated Task)** | **Name in UniProtKB** | **Number of Clusters in UniRef50** | **Average Similarity** | **Number of Clusters with >40% Similarity** |
|---|---|---|---|---|
| rpoA (RIF) | DNA-directed RNA polymerase subunit alpha | 1772 | 0.29 | 293 |
| rpoB_rpoC (RIF) | DNA-directed RNA polymerase subunit beta | 4008 | 0.20 | 657 |
| Rv2752c (RIF) | Multidrug efflux pump Tap | 3 | 0.28 | 236 |
| pncA (PZA) | Nicotinamidase pyrazinamidase | 355 | 0.33 | 60 |
| clpC1 (PZA) | ATP-dependent Clp protease ATP-binding subunit ClpC1 | 52 | 0.20 | 2 |
| panD (PZA) | Aspartate 1-decarboxylase | 666 | 0.34 | 230 |
| Rv1258c (PZA) | Multidrug efflux pump Tap | 3 | 0.44 | 1 |
| PPE35 (PZA) | PPE family protein PPE35 | 1 | 0.10 | 0 |
| Rv3236c (PZA) | Probable conserved integral membrane transport protein | 0 | N/A | N/A |
| mshA (INH) | D-inositol 3-phosphate glycosyltransferase | 3776 | 0.28 | 11 |
| Rv1258c (INH) | Multidrug efflux pump Tap | 3 | 0.44 | 1 |
| fabG1 (INH) | 3-oxoacyl-acyl-carrier-protein reductase MabA | 2 | 0.32 | 3 |
| inhA (INH) | Enoyl-[acyl-carrier-protein] reductase [NADH] | 535 | 0.22 | 0 |
| furA (INH) | Transcriptional regulator FurA | 18 | 0.46 | 11 |
| katG (INH) | Catalase-peroxidase | 717 | 0.22 | 90 |
| ndh (INH) | Type II NADH:quinone oxidoreductase Ndh | 1 | 1.0 | 1 |
| ahpC (INH) | Alkyl hydroperoxide reductase C | 4460 | 0.23 | 20 |
| Rv2752c (INH) | Ribonuclease J | 1680 | 0.28 | 236 |
| S (SARS-CoV-2) | Spike Glycoprotein | 171 | 0.20 | 0 |
| GFP (Aequorea victoria) | Green fluorescent protein | 46 | 0.33 | 9 |

**Table S3:** Information on cluster selection in Uniref50[8] for RIF, PZA, INH, SARS-CoV-2, and GFP datasets. The table displays: (1) the gene name from UniProtKB used to search for similar clusters in UniRef50, (2) the number of clusters found in UniRef50 with that gene name, (3) the average sequence similarity of representative sequences from all clusters to the reference gene sequence, and (4) the number of clusters with over 40% sequence similarity to the reference gene sequence.

| **ProteinGym Dataset Name** | **Name in UniProtKB** | **Number of Clusters in UniRef50** | **Average Similarity** | **Number of Clusters with >40% Similarity** |
|---|---|---|---|---|
| AACC1-PSEAI-Dandage-2018 | Gentamicin 3-N-acetyltransferase; AAC(3)-I family aminoglycoside N-acetyltransferase; acetyltransferase | 305055 | 0.23 | 34 |
| A4GRB6-PSEAI-Chen-2020 | VIM-2 metallo-beta-lactamase;Anti-Pycsar protein Apyc1, Beta-lactamase | 76291 | 0.24 | 37 |
| A4D664-9INFA-Soh-CCL141-2019 | Polymerase basic protein 2 | 68 | 0.17 | 0 |
| A4-HUMAN-Seuma-2021 | Amyloid-beta precursor protein;Amyloid-beta | 754 | 0.22 | 30 |
| A0A140D2T1-ZIKV-Sourisseau-growth-2019 | Genome polyprotein | 2752 | 0.12 | 13 |

**Table S4:** Information on cluster selection in Uniref50[8] for ProteinGym datasets. The table displays: (1) the gene name from UniProtKB used to search for similar clusters in UniRef50, (2) the number of clusters found in UniRef50 with that gene name, (3) the average sequence similarity of representative sequences from all clusters to the reference gene sequence, and (4) the number of clusters with over 40% sequence similarity to the reference gene sequence. Multiple gene names used for searches are separated by a semicolon.
